# Mapping global marine biodiversity under sparse data conditions

**DOI:** 10.1101/2023.02.28.530497

**Authors:** Damiano Righetti, Meike Vogt, Nicolas Gruber, Niklaus E. Zimmermann

## Abstract

Sparse and spatiotemporally highly uneven sampling efforts pose major challenges to obtaining accurate species and biodiversity distributions. Here, we demonstrate how limited surveys can be integrated with global models to uncover hotspots and distributions of marine biodiversity. We test the skill of recent and advanced species distribution model setups to predict the global biodiversity of >560 phytoplankton species from 183,000 samples. Recent setups attain quasi-null skill, while models optimized for sparse data explain up to 91% of directly observed species richness variations. Using a refined spatial cross-validation approach to address data sparsity at multiple temporal resolutions we find that background choices are the most critical step. Predictor variables selected from broad sets of drivers and tuned for each species individually improve the models’ ability in identifying richness hotspots and latitude gradients. Optimal setups identify tropical hotspots, while common ones lead to polar hotspots disjunct from general marine diversity. Our results show that unless great care is taken to validate models, conservation areas in the ocean may be misplaced. Yet a game-changing advance in mapping diversity can be achieved by addressing data-sparse conditions that prevail for >80% of extant marine species.

**Authorship statement:** All authors designed the research and contributed to the writing. D.R. designed the multiscale validation and predictor selection methods, developed the figures with input by M.V. and N.E.Z., performed research, and wrote the first draft.

## Introduction

Accurate biodiversity maps are essential for ecological assessments and inference, the design of reserves, and protection of natural assets. Such maps critically depend on biological occurrence data collected through traditional and molecular field surveys, and the quantities of these data are surging (Heberling *et al*. 2021). Such data have helped demonstrate the vital role that biodiversity plays for ecosystem services, such as productivity, carbon storage (Poorter *et al*. 2015; Liang *et al*. 2016) and resilience to climate extremes (Isbell *et al*. 2015; Duffy *et al*. 2016). Nevertheless, spatial patterns of diversity have been unknown for over 80% of species (Mora *et al*. 2011), including many protists, fungi, insects, animals (Sogin *et al*. 2006; Kass *et al*. 2022) and functionally rich marine microbes (Paoli *et al*. 2022).

Until now, a cardinal problem with the global biodiversity descriptions of these groups have been data limitations, including spatiotemporal sampling biases (Hortal *et al*. 2015; Meyer *et al*. 2016), varying taxonomic coverage or quality of occurrence records (Selig *et al*. 2014), and massive data gaps (Robinson *et al*. 2011; Meyer *et al*. 2015). These biases and gaps have been particularly acute in open ocean systems (Robinson *et al*. 2011; Menegotto & Rangel 2018). Besides affecting biodiversity assessments (Chaudhary *et al*. 2017), these issues induce considerable uncertainty in determining the key environmental factors defining a species’ niche (Beaugrand *et al*. 2013 vs. Rivero-Calle *et al*. 2015). This latter issue affects the skill of statistical predictions (Araújo *et al*. 2019; Santini *et al*. 2021) with a type of mapping tool widely used in biodiversity and conservation science: species distribution models (SDMs).

SDMs are standard tools for mapping biodiversity from occurrence records (e.g., Benedetti *et al*. 2018; Zurell *et al*. 2018; Araújo *et al*. 2019; Carlson *et al*. 2022). By resolving the wealth of species at a particular location from the overlap of species’ distributions, these models open paths for mapping taxonomic and functional (e.g., trait or metabolic) diversity. SDMs rely on accurate estimates of a species’ niche (*sensu* Hutchinson 1957) derived from empirical relationships that exist between species’ occurrences and environmental factors (Guisan & Zimmermann 2000; Lee-Yaw *et al*. 2016). These relationships are projected in space, assuming a lack of dispersal barriers, as is often the case with oceanic taxa (Cermeño and Falkowski 2009, Whittaker and Rynearson 2017; but see Ward et al. 2021). Since the 1990s, at least 6,500 SDM studies have been conducted (Araújo *et al*. 2019). However, only about 400 of these studies have addressed marine taxa (Melo-Merino *et al*. 2020), and fewer than 15 have reported on open-ocean diversity globally, examining fish (Worm 2005; Cheung *et al*. 2009; Jones & Cheung 2015), mammals (Kaschner *et al*. 2011; Auber *et al*. 2022), brittle stars (Woolley *et al*. 2016), plankton (Thomas *et al*. 2012; Ladau *et al*. 2013; O’Brien *et al*. 2016; Jorda *et al*. 2019; Righetti *et al*. 2019; Benedetti *et al*. 2021) and other taxa (Selig *et al*. 2014; García Molinos *et al*. 2016). Marine SDM applications gained momentum since 2005 (Melo-Merino *et al*. 2020), but a summary of 236 studies has shown that data uncertainties are rarely addressed (Robinson *et al*. 2017). This contrasts with the fact that sampling biases are omnipresent in oceans (Menegotto & Rangel 2018), which likely create a major source of model errors (Bardon et al. 2021). Though model choices are critical to address the biases (Phillips *et al*. 2009; Righetti *et al*. 2019; Benedetti *et al*. 2021; Auber *et al*. 2022) the extent to which model choices can overcome data limitations has not been established globally.

Phytoplankton represent an abundant and diverse form of marine life (de Vargas *et al*. 2015). These microbes set a powerful precedent to test the skill of SDMs, as their data feature at least five challenges (refer to *SI*, Box 1 for details). Notably, (1) data points are sparse for most species (Righetti *et al*. 2019); (2) sampling is spatiotemporally biased (Righetti *et al*. 2020); (3) many samples suffer from incomplete species detection (Cermeño *et al*. 2014; Rodríguez-Ramos *et al*. 2015); (4) detection probability is low owing to seasonal dynamics in species occurrence (Righetti *et al*. 2019); and (5) absence data for most species are unavailable (Barton *et al*. 2016; Righetti *et al*. 2020). Overcoming these challenges requires careful choices regarding the setup of SDMs, i.e., those associated with the selection of background data (pseudo-absences), predictors, and thresholds to infer species’ presence and absence. In addition, the lack of reliable absence data leads to a validation problem, as skill tests typically combine the rates of both correctly predicted presences and absences (Allouche *et al*. 2006).

To advance biodiversity models suitable for these major data challenges, we use 182,303 pelagic samples spanning 567 phytoplankton species (Righetti *et al*. 2019) to calibrate 18 SDM setups that span recently used to more sophisticated/new background, predictor, and thresholding choices. We address (1) the spatiotemporal sampling bias and missing absence records, by selecting background data via three methods: the first selects data from the full ocean at random, while the refined methods select data from sampling sites of the general target group studied (i.e., phytoplankton) or specific taxa (e.g., diatoms), mirroring sampling biases (Phillips *et al*. 2009). Next, we address (2) incomplete knowledge of the factors delimiting a species’ niche by using an invariant set of predictors and two methods to select predictor sets and ensembles optimized for each species, accounting for species’ possible idiosyncratic dependencies on factors. Finally, we revisit (3) the paradigm that probabilistic SDM projections, summed across species, explain species richness more accurately than presence-absence (0/1) projections (Calabrese *et al*. 2014). This view stems largely from terrestrial cases where 0/1 projections overpredict species numbers (Dubuis *et al*. 2011; Guisan & Rahbek 2011; Guillera-Arroita *et al*. 2015; Zurell *et al*. 2020). However, direct marine tests have not been attempted, as the existing work focuses on thresholds optimizing the convergence between both approaches, rather than the 0/1 approach (Benedetti *et al*. 2021). To date, the ability by which either projection type can predict global biodiversity variations, including pervasive slopes from poles to tropics, is an open question.

To test to what degree SDMs translate into accurate biodiversity distributions we first build a method to extract richness test data. We then use spatial-block cross validation (sbCV) (Roberts *et al*. 2017; Ploton *et al*. 2020) to test predictions in a spatiotemporally exact manner against the richness found in raw data at over 20 space-time resolutions (Methods). By extracting global test data from presence only data, our sbCV bypasses the need for absences. While the models are trained using monthly data, the CV extends to multi-month test scales (Araújo *et al*. 2019) important for marine microbes whose detectable ranges may shift seasonally. Using 1 million phytoplankton records, we find that optimized background, predictor and threshold choices drastically improve the global mapping of plankton diversity.

## Methods

### Basic analysis setup

We employed 18 SDM setups (Table 1) with alternating choices of background data, predictor variables, and projection types. These setups were used to fit three SDM algorithms: generalized linear models (GLMs), generalized additive models (GAMs), and random forests (RFs). We summarize our workflow of building SDMs in Figure 1. Each SDM was subjected to random fourfold CV (step 6), and successful SDMs were combined to ensembles (step 7). The models were used to predict richness at monthly to annual timescales (step 8) and evaluated using global sbCV (step 9). sbCV scores were fed into an ANOVA to quantify effects of choices on SDM skill. All steps were run in R v.3.6.3.

**Fig. 1.**
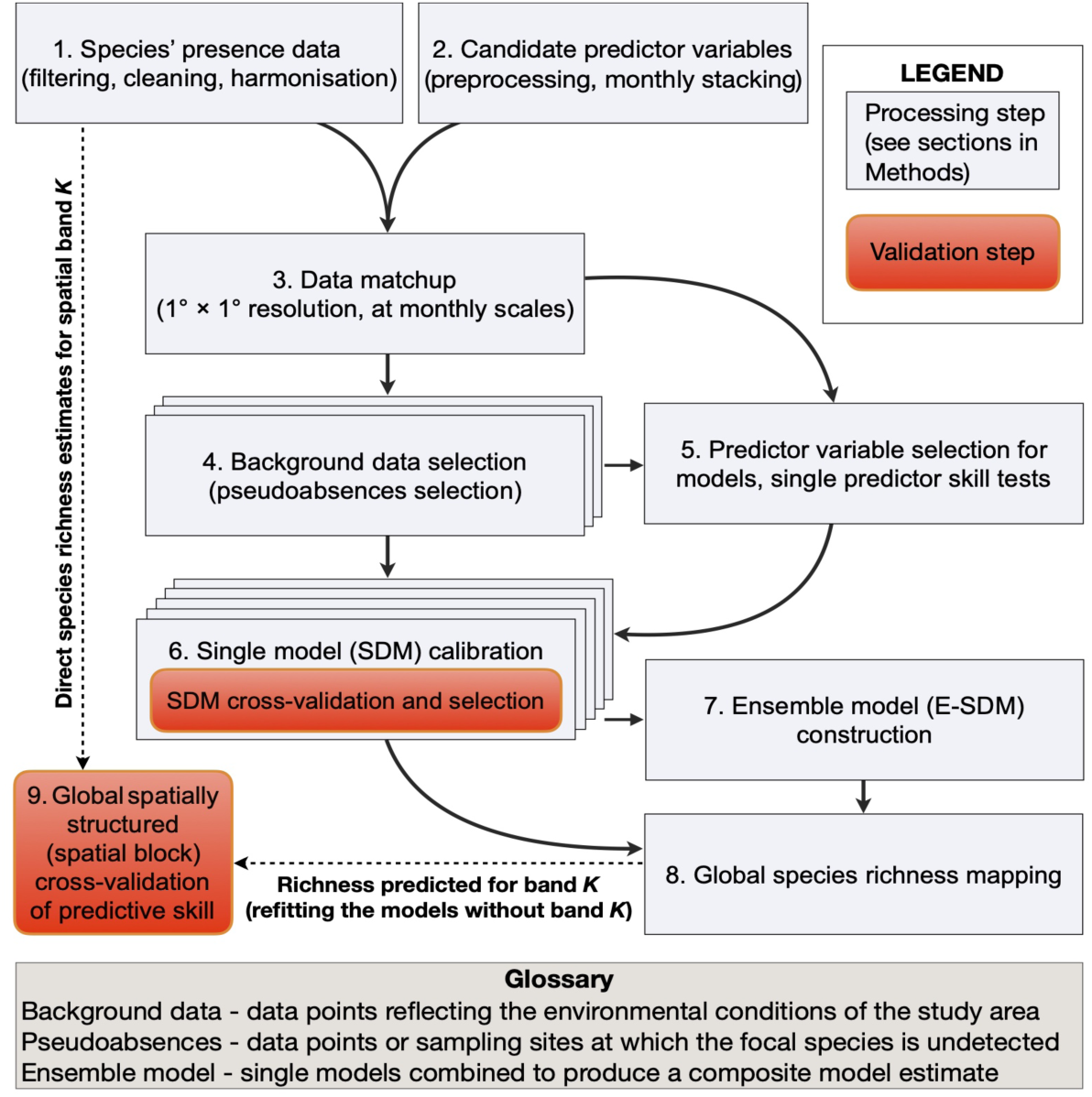
Flow diagram of model building steps. The assembly and validation of species distribution models (SDMs) consists of nine steps, including SDM calibration, individual-model random fourfold cross validation, ensemble model generation, predictions of species richness, and their global spatial-block cross validation.

**Table 1.** Methodological setup. SDMs used to predict diversity from biased and sparse data efforts employed varying background (3), predictor (3), and projection (2) choices, yielding 18 setups used to fit 3 algorithms. Setups 10 to 18 used probability-of-presence (PRB), and setups 1 to 9 used presence-absence (PA) projections, all other settings are identical.

| Setup no. | Projection type choice | Predictor choice | Background choice | Spp. modeled (considered)<sup>§</sup> |
| --- | --- | --- | --- | --- |
| 1 (10) | PA (PRB) | ensemble | group-specific target group | 536<sub>GAM</sub>, 529<sub>GLM</sub>, 538<sub>RF</sub> (566) |
| 2 (11) | PA (PRB) | top member | group-specific target group | equal to S1 |
| 3 (12) | PA (PRB) | generic | group-specific target group | 402<sub>GAM</sub>, 389<sub>GLM</sub>, 415<sub>RF</sub> (447)<sup>§</sup> |
| 4 (13) | PA (PRB) | ensemble | total target group | 548<sub>GAM</sub>, 543<sub>GLM</sub>, 551<sub>RF</sub> (566) |
| 5 (14) | PA (PRB) | top member | total target group | equal to S4 |
| 6 (15) | PA (PRB) | generic | total target group | 409<sub>GAM</sub>, 406<sub>GLM</sub>, 427<sub>RF</sub> (447)<sup>§</sup> |
| 7 (16) | PA (PRB) | ensemble | full ocean | 567<sub>GAM</sub>, 566<sub>GLM</sub>, 566<sub>GLM</sub> (567) |
| 8 (17) | PA (PRB) | top member | full ocean | equal to S7 |
| 9 (18) | PA (PRB) | generic | full ocean | 438<sub>GAM</sub>, 431<sub>GLM</sub>, 447<sub>RF</sub> (447)<sup>§</sup> |

<sup>\*</sup>"generic" predictors refers to a fixed set of predictors for all species: T, pCO<sub>2</sub>, sea surface wind stress, and N<sup>\*</sup>
<sup>§</sup>The species considered for modeling had ≥ 24 presences. Only 402 to 447 species received ≥ 24 presences when using fixed predictors, as pCO<sub>2</sub> fields had a limited spatial coverage. Species considered for richness analyses herein scored a TSS ≥ 0.35 in fourfold random CV

#### 1. Phytoplankton data

We compiled 1,056,363 species presence records sampled near the ocean surface (mean depth = 5.4 m, SD = 6.9 m) for modeling and direct analyses (Fig. 1; step 1). Data and cleaning protocols are accessible through PhytoBase (Righetti *et al*. 2020), selecting for download year 2015 and excluding species without any records within the monthly mixed layer (de Boyer Montégut 2004; temperature criterion), yielding 1300 species (482 *Bacillariophyceae*, 643 *Dinophyceae*, 127 *Haptophyta*, 4 *Cyanobacteria*). Data originated from GBIF (www.gbif.org), OBIS (www.obis.org), Buitenhuis et al. (2013), Villar et al. (2015), and Sal et al. (2013). We restricted data to seas of ≥ 200 m depth (Amante & Eakins 2009) and surface salinity ≥ 20 (Zweng *et al*. 2013), excluding ecologically more complex nearshore seas.

#### 2. Candidate predictors

26 oceanographic predictor variables, detailed in Table S1, were gridded at 1×1° (WGS84) and monthly (*n* = 12) climatological resolution (Fig. 1; step 2). The physical, chemical, and biological predictors represent key facets of phytoplankton niches affecting the species’ physiology, growth and competition (Margalef 1978; Litchman *et al*. 2007; Boyd *et al*. 2010; Brun *et al*. 2015; Dutkiewicz *et al*. 2020; Seifert *et al*. 2020).

#### 3. Data matchup and species selection

For each species, presence data were binned to 1×1° cells at monthly resolution, matching predictor data resolution, yielding 245,322 binned presences (Fig. 1; step 3). The latter cover 17,333 (3.5%) of the ocean’s monthly 1° cells (see fig. S1). The binning of multiple samples into 1° cells per calendarial month ameliorates the low species detection power of single samples (Cermeño *et al*. 2014), but neglects potential interannual changes in species distribution. We expect no major loss in SDM accuracy, since global environmental gradients exceed possible environmental changes from 1950 to 2010 by far, over which 96.1% of our presences were collected. We set the minimum number of presences to 24 for inclusion of species into SDMs, yielding 402–567 species (Table 1) across SDM setups (247 *Bacillariophyceae*, 268 *Dinophyceae,* 38 *Haptophyta*, 4 *Cyanobacteria*, 10 others).

#### 4. Background selection

We selected background data (Fig. 1; step 4) at a ratio ten times higher than presence data available for each species (Barbet-Massin *et al*. 2012). Background data were selected from open ocean cells at monthly 1° resolution, via three methods. Two of these methods correct for original survey bias by introducing a similar background-data distribution bias (Phillips *et al*. 2009). We also tested spatial thinning as a means to reduce bias (*SI*, section S1), yet we discarded this method due to substantial data loss.

1. *<u>Random</u>*: In line with recently used plankton SDMs (e.g., Pinkernell and Beszteri 2014, Brun et al. 2015, Barton et al. 2016, Jensen et al. 2017), we selected background data spatially and temporally at random from the full study area, i.e. monthly 1° cells of the global ocean.
2. *<u>Total target group</u>*: We next selected background data from the subset of monthly 1° cells that reported presences (see fig. S1A) of any of the 1300 species in our original dataset (step 1). We expect the density of cells spanned by this “total target group” (TTG) to mirror the presence survey effort and bias applied to our study species (Phillips et al. 2009).
3. *<u>Group-specific target groups</u>*: Since many surveys report only occurrences of specific plankton taxa (Righetti *et al*. 2020), we also defined group-specific target groups (GSTGs) separately for *Bacillariophyceae*, *Dinoflagellata*, and *Haptophyta*. These major groups cover 96% of the species in our dataset. All species, except the *Bacillariophyceae*, served as GSTG for any remaining data-deficient group, such as small-sized *Cyanobacteria*. We omitted *Bacillariophyceae* from the target-group definition of the remaining taxa because they feature a heavy data cluster in the North Atlantic (Righetti *et al*. 2020) from net-sampling methods that neglect small-sized plankton (Richardson *et al*. 2006).

Within approach 2–3, we sampled background data in an environmentally stratified manner, dividing both the monthly T and MLD gradient spanned by the target group’s presences into nine regular intervals, yielding 81 strata (T × MLD combinations). These two factors were used as they are ecologically highly relevant (Margalef 1978; Brun *et al*. 2015) but weakly correlated (Table S2). In each stratum, data were selected at random from and in proportion to the 1° cells containing presences of the target group. Thus, the quantity and distribution of monthly 1° cells sampled (rather than the quantity of species records that is affected by species richness) in the target group served as proxy for original sampling effort.

#### 5. Predictor selection

Based on an initial ranking of 26 candidate predictors (Table S1) in discriminating species’ presences vs. background data, we selected the likely key drivers and most skillful predictors for SDMs (Fig. 1; step 5). We fitted single-variable GLMs, GAMs, and RFs to the presence and GSTG background data of each species (see *SI*, section S2) and assessed model fit by the adjusted *R*^2^ for GLMs and GAMs and the out-of-bag error statistic for RFs. We ranked the candidates per species using the mean of the GLM, GAM, and RF based rankings of model fits for the final ranking per species. From this, we modeled each species by three ways of predictor choice. In all cases, we selected predictors such that they were not highly correlated (i.e., |*ρ*| ≤ 0.7; Table S2) within SDMs (Dormann *et al*. 2013).

1. *<u>Generic predictors</u>:* We first used a fixed set of the same four predictors across all SDMs. The four selected predictors are the potential key dimensions of phytoplankton niches (see *SI*, section S3) and show particularly low pairwise correlations, serving to maximize joint explanatory power. Namely, we selected: sea surface temperature, wind stress, pCO_2_, and N*.
2. *<u>Specific-specific ensembles of predictors</u>:* To capture possible distinctions between species’ niche dimensions, we next generated five sets of four predictors individually optimized for each species to fit five member SDMs. For each of the sets, we randomly selected four predictors correlated at |*ρ*| ≤ 0.7 from the species’ top 10 ranked candidate predictors. To avoid overrepresentation of individual predictors per species, we selected each predictor (including log10 versions) only up to twice across the five sets. In cases were the top-ranked candidates provided insufficient predictors to define five sets, candidates ranked >10 (up to 26) were selected. Finally, we built an ensemble SDM by unweighted averaging of those member SDMs, which scored a TSS ≥ 0.35 (using fourfold CV, step 6). This allowed covering a large set of relevant predictors while keeping predictor numbers low per SDM (Breiner et al. 2015).
3. *<u>Species-specific top predictors</u>:* Out of the five member SDMs fitted by the five sets, we finally used the member per species with the highest TSS score (at least 0.35, fourfold CV.

#### 6. SDM calibration and fourfold CV

The predictors were used to fit three SDM algorithms (Fig 1; step 6) that provide varying degrees of flexibility in statistical response curves: GLMs (R package *stats*), GAMs (*mgcv*; Wood, 2001) and RFs (*randomForest*; Liaw & Wiener 2002). We used four predictors per SDM and we tuned each algorithm to fit rather simple curves, given the sparse data (Merow *et al*. 2014) with 50% of our species having <160 presences (25% <54 presences). GAMs included smoothing terms with five basis dimensions, estimated with penalized regression splines, without penalization to zero for single variables. GLMs used linear and quadratic terms and stepwise bidirectional predictor selection. RFs used simple terms, 4,000 trees, and single end-node size. Presences and pseudo-absences (i.e., background data) were weighted equally in GLMs and GAMs to obtain prevalence weights of 0.5. RFs subsampled background data equal to the number of presences. We tested SDM skill by fourfold CV. We selected 75% (three folds) of the data at random to train SDMs, keeping prevalence in data subsets equal to that in total data, predicting to the withheld 25% (one fold) iteratively, until each fold part was used once for testing, and three times for training. We used the true skill statistic (TSS) as test metric (Allouche *et al*. 2006). We applied TSS-maximizing thresholds to convert probabilistic SDM raw projections (or ensembles of SDMs) into presence-absence (PA) projections required by this test. We retained SDMs with a TSS ≥ 0.35 for richness analyses, resulting in 402–567 species per setup (Table 1).

#### 7. SDM projection and ensemble building

Successful SDMs (TSS ≥ 0.35) were projected onto the monthly environmental data fields (see step 2) to obtain probabilistic (PRB) projections of species’ presence at monthly 1° resolution. PRB projections were converted into presence–absence projections by using TSS-maximizing thresholds (*presenceAbsence*, Freeman & Moisen 2008). For each species, we derived an ensemble model (E-SDM) projection (Fig. 1; step 7), using the mean of the monthly PA projections of successful members (*n* = 1–5) and 0.5 as the final threshold per monthly 1° cell. PRB E-SDM projections were obtained by using the mean of PRB projections, without final thresholding.

#### 8. Global diversity maps

For each SDM setup, we used the sum of the stacked PA or PRB projections of the species to predict species richness at monthly 1° resolution (Fig. 1; step 8). Multi-monthly estimates were derived by integrating the PA projections of month *n* + *m* (consecutive) months. Maps visualize the means of the twelve estimates throughout the year.

#### 9. Spatial block CV (sbCV)

We used sbCV (Fig. 1; step 9) to assess the skill of our SDMs to generalize species richness globally to areas withheld from SDM training, while separating training and test data spatially (Ploton *et al*. 2020). This CV involved two subsequent steps:

1. *<u>Test-data determination</u>.* We repeatedly selected predefined numbers of samples from spatial bands of 15° latitude width (Fig. 2; *SI*, sections S4–5). Determining the integral species richness from samples, pooled per latitude band served to estimate richness. This pooling, in part, reduces the strong under-sampling of richness by traditionally small seawater volumes (Cermeño et al., 2014). We took exact latitude-longitude-year-month-day-depth IDs in the raw presences, discarding NA depths, to define 182,303 samples (3,909 discarded). We accepted 63,441 samples, each containing ≥ 5 species (refer to fig. S2, and *SI* section S5) for richness analyses at 30 (*n* samples × *m* cells) aggregation scales (Fig. 2B). Per band, we selected two samples (up to ten) from up to six monthly 1° cells selected at random. Richness was determined for species that were successfully modeled by the SDM setup (Table 1) tested. Out of 30 (5 × 6) analysis scales, 23 found sufficient samples in at least 9 bands. We randomly selected samples for each samples × cells combination 1,000 times, to obtain 23 richness mean trends (“observed truths”) robust to data noise (Fig. 2C).
2. *<u>Test</u>.* We refitted the SDMs built during steps 1–7 based on *K*−1 latitude bands (Fig. 2A), We left model setups unchanged, except for updating presences to background weights (prevalence 0.5), until each band was withheld once from SDM training, and received a prediction. For each of the 23 scales, we contrasted the observed truth to the richness predicted for exact same spatiotemporally explicit 1° cells (*SI*, section S6). We ran 3.93 million predictions for the sbCV (3 algorithms × 18 setups × *n* species by setup × 12 bands × 12 months), yielding 1,242 predicted-to-observed richness contrasts (Fig. 2D–E; 3 algorithms × 18 setups × 23 scales). The contrasts were built to quantify similarity between observed and predicted species richness. We used the Spearman’s *ρ* of each contrast as our main CV score. Pearson’s *r*, mean absolute error (MAE), and root mean square error (RMSE) served for robustness tests (*SI*, section S7). The latter two were run with raw and modelled data scaled to 1 at maxima. This is required because raw data are too sparse to represent realistic absolute numbers. Yet, the raw data are sufficient to indicate global latitudinal distribution patterns of diversity at manifold data resolutions, which we test here.

**Fig. 2.**
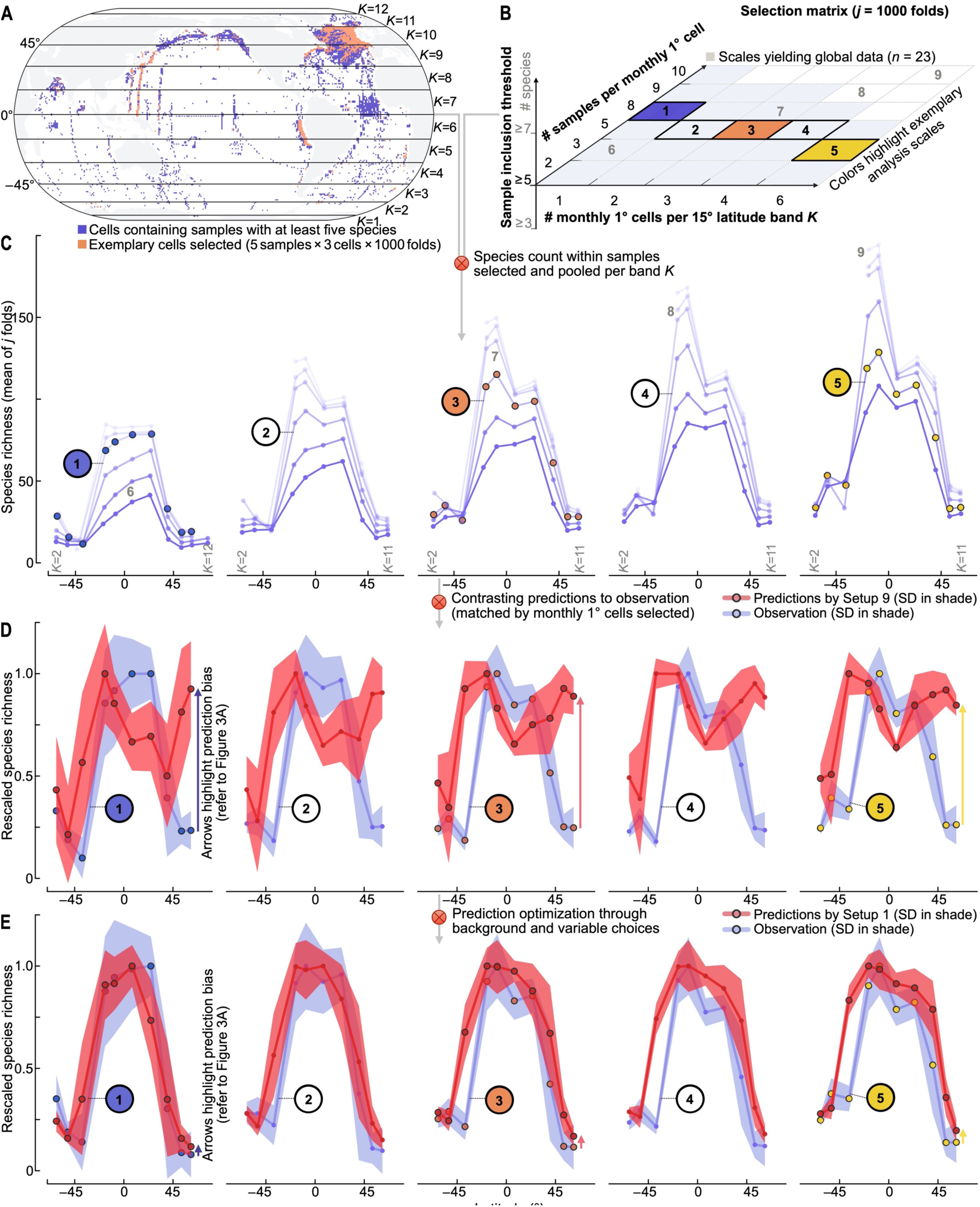
Latitudinal richness gradients deduced from sparse global data. (A) Locations of 1° cells containing samples. (B) Matrix defining the number of samples selected and pooled at varying spatial or temporal extents (subsampling design) from each of twelve latitude bands. Specifically, the temporal integration of diversity examined broadens with increasing numbers of monthly 1° cells selected (*N* = 1–6). (C) Latitudinal species richness gradients resulting at 30 scales of analysis defined in panel (B). Blue dots (connected by lines) show the means of applying the matrix in (B) 1000 times. (D, E) The predicted richness (red) is contrasted to the matching observed richness (blue) at five scales of analysis using once a baseline SDM setup (D) and once an advanced setup (E). The numbers in panels C–E highlight results at specific extents of analysis, defined in in the resampling matrix (panel B).

### Richness slope comparisons

As an additional skill test chiefly robust to residual biases in our re-sampling, we derived the linear fits of the predicted (sbCV) and observed (test data) richness along global temperature or latitude and contrasted predicted with observed slopes.

### Statistical effect-size analysis

To quantify the effect of different choices on SDM skill, we used ANOVA (*stats* package) based on linear models, with CV-score as the response, and background, predictor, projection, algorithm choice, and linear interactions, as factors.

## Results

### Latitudinal variations

Our test-data generation method for addressing biases in survey effort at the sample, cell, and latitude band levels reveals the sensitivity global latitudinal gradients of phytoplankton richness found at monthly to multi-monthly scales. By latitude, maxima occur near the equator, reaching ∼35 to 160 species when using two to 30 samples per band (Fig. 2A–B), but in the subtropics at multi-monthly timescales (Fig. 2C). All diversity gradients show a clear tropical-to-polar decline that wanes or inflects at ∼45° N/S.

This multi-faceted empirical baseline of global species richness (i.e., latitudinal variation found in samples pooled at different spatial and temporal extents) is reproduced by our different model setups with widely varying accuracies: the setups explain on average between 10.9% and 92.7% (mean *r^2^*) of the 23 latitudinal richness variations observed. Setups with generic predictor and random background choice (S9, S18) are inaccurate (Fig. 3A), failing to reproduce the sign of the latitudinal gradient, specifically at northern latitudes (Fig. 2D). The refined setups, however, closely match the variations observed (Fig. 2E, fig. S3).

**Fig. 3.**
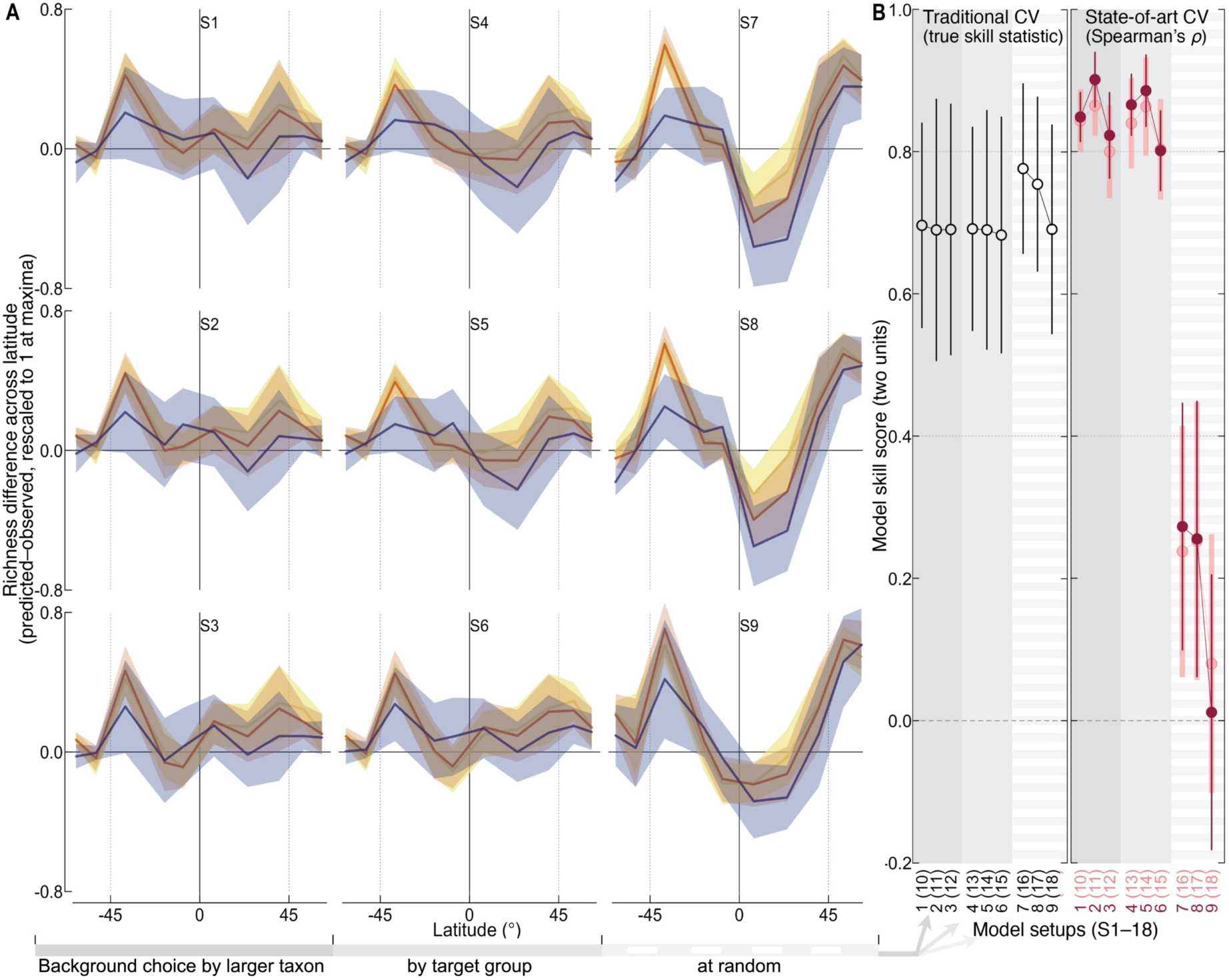
Bias and skill of all model setups (SDMs) in predicting species richness globally on multiple timescales. (A) Prediction bias across latitudes, expressed as predicted minus observed phytoplankton species richness (both rescaled to 1 at maxima). Colors denote three exemplary timescales of analysis (monthly, tri-monthly, six-monthly; see Fig. 2B) using the richness predicted by SDM setups 1–9 under the generalized additive model (GAM) algorithm. Lines show means of 1000 folds (SD in shading), with larger deviation from the *x*-axis indicating stronger error. (B) Spatial block cross validation (CV) scores of all 18 SDM setups under the GAM algorithm (dots are means, lines ± 1 SD). (C) Random split-data CV scores (dots are means, lines ± 1 SD). Results are sorted by background choice.

Similarly, the skill of SDM setups in globally predicting species richness across bands withheld from the SDM training steps is varied (Fig. 3B, state-of-art CV), ranging from essentially no skill (mean *r^2^* of 0.05%, mean *ρ* of 0.01) for the random background data choice (SDM setups 7–9, 16–18) to highly faithful predictions that explain up to 91.4% (*r*^2^, mean *ρ* of 0.92; Table S3) of the latitudinal richness variations observed (setups 1–2, 10–11).

### Determinants of predictive skill. *Background choice*

In our system, background choices exert the dominant effect on model skill, exceeding the effect size or uncertainty induced by SDM algorithm choice by 3.9 to 21.5-fold (fig. S4). SDMs calibrated using the random background result in predictive failure (Fig. 3B, state-of-art CV, and Table S3) at a mean *ρ* below 0.4 and *r*^2^ < 0.25 (sbCV), irrespective of other calibration choices and they reveal highly skewed global patterns in predicted-to-observed differences across latitude (Fig. 3A). Using refined background data sampled from the total target group (TTG) or from within major taxa (GSTG choice) results in a large gain in model skill (Fig. 3B, state-of-art CV), with all setups explaining over 75% (mean *r*^2^, mean *ρ ≥* 0.76) of observed variations. Target-group background choices lead to highly significant skill gains in each sbCV metric compared to random background choice (paired Wilcoxon test, *p* <0.001) for each statistical SDM algorithm (fig. S5).

Different background choices result in radically different patterns of phytoplankton species richness (Fig. 4 A–B vs. C–D). Only background choices designed to filter original survey bias (Fig. 4C–D) led to spatiotemporal species richness patterns that replicate the equator-to-pole declines found in the observational data (Fig. 2; Fig. 4 dots).

**Fig. 4.**
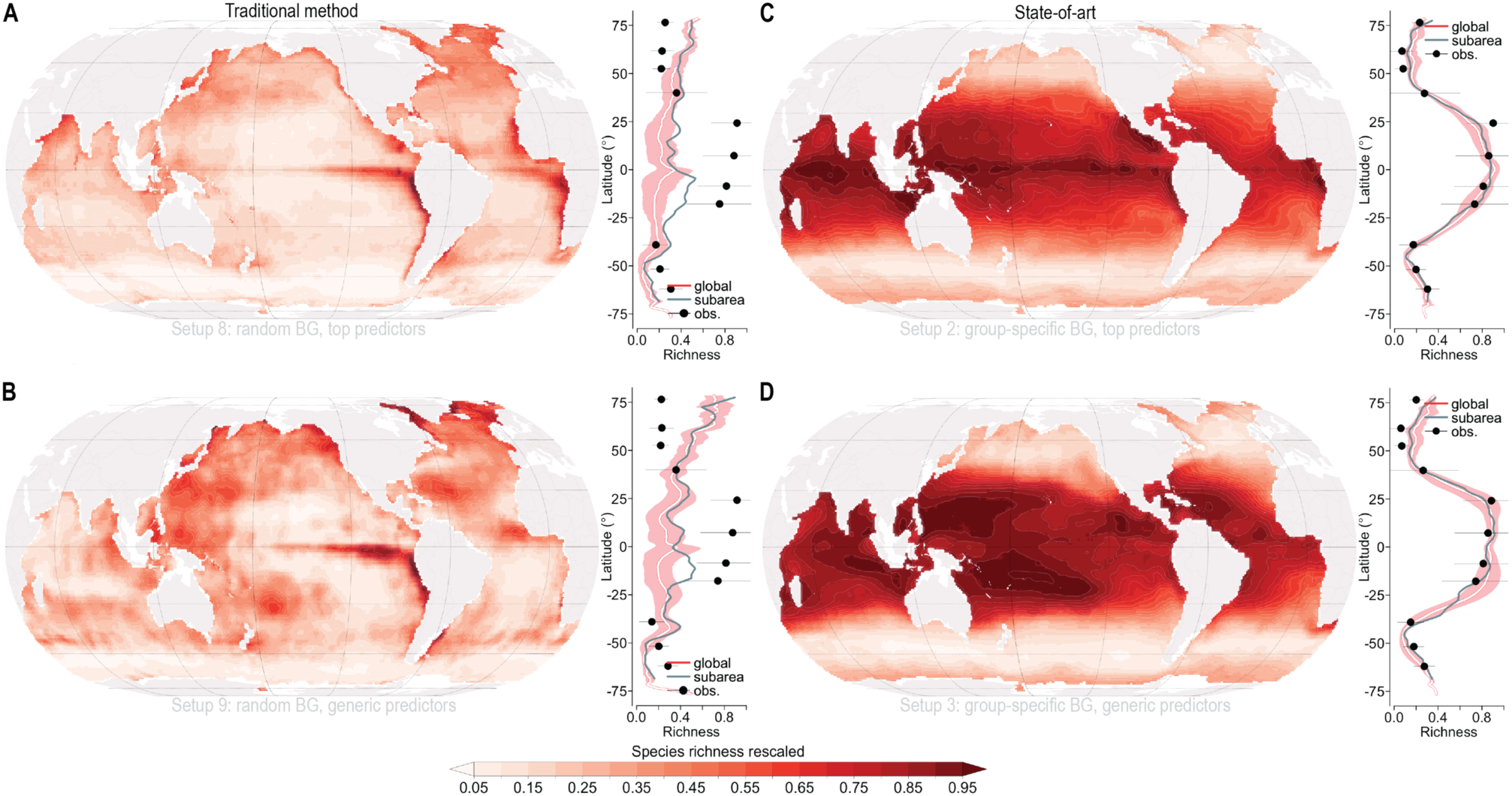
Global distributions of phytoplankton species richness at monthly resolution predicted by model setups. Maps show the annual mean of monthly species richness predicted by models with random background choice (A–B) as contrasted to setups with newly refined target-group backgrounds (C–D), using generalized additive models. White and green lines indicate the means per degree latitude for the full ocean area (white, SD in red shading) and the subarea in which samples were available (green, see Fig 2A). Dots show the mean richness found in pooled samples (lines for SD) using 1000 selection folds (Methods). Richness maxima were rescaled to 1 for comparisons.

The effect of target-group (GSTG and TTG) background choice is also striking in the determination of global relationships between species richness and sea surface temperature (fig. S6) or latitude (fig. S7). Global linear slopes of ln (species richness) across sea surface temperature, determined at the 23 resolutions of sampling-data analysis, are best replicated by setup 1 (using GSTG), followed by setup 4 (using TTG), with robust outcomes across the 23 data resolutions and three SDM algorithms tested (fig. S6). Across latitude, the model setup 1 performs best, followed by setup 2 (fig. S7).

While SDM setups 1–3 (10–11) with random background choice reach the lowest skill scores based on state-of-art spatial block CV (Fig. 3B) and the latitudinal and thermal slope analysis (fig. S6–7), they reach the highest skill scores as assessed by the widely used random CV approach (Fig. 3B, and fig. S5).

#### Predictor variable choices

Our refined predictor choices, which address modeled species individually rather than all species in a common way, had important second-order effects on predictive skill, leading to skill gains on top of those induced by optimal background choices.

The effect size of predictor choices on SDM skill as assessed by Spearman’s *ρ* (sbCV) was about that of algorithm choice (fig. S4) but lower for other CV metrics (MAE, RMSE, *r*). Species-specific predictor variables choices improved SDM skill significantly relative to the generic predictor choice as assessed by *ρ* (paired Wilcoxon test, *p* <0.001 and *n* = 138 for each algorithm except RF, ns), but results remained more mixed for alternative CV metrics (*SI*, supplementary results).

The effects of predictor choice on mapped richness distributions are substantial. Patchiness in spatial richness patterns is found based on generic predictor selection (Fig. 4B) but clearly reduced as a consequence of species-specific predictor selection (Fig. 4A). Using the most refined (GSTG) background choice, richness maxima shift within the Pacific as a function of the predictor choice (Fig. 4C–D). Basin-scale hotspots of species richness at monthly time resolution emerge near the equator based on species-specific predictors, while generic predictors predict hotspots at ∼20° N/S (Fig. 5C, red vs. yellow). Similarly, sensitivity to predictor choice is found for subgroups with even sparser data, such as the haptophytes (Fig. 5B, D), while richness coldspots are generally more stable under differing predictor choices. Diversity features, particularly pantropical richness hotspots, are altered depending on the temporal resolution of the analysis (Fig. S9). At monthly to annual resolution scales, subtropical and midlatitude richness maxima are more common (fig. S9) than at the monthly scale (Fig. 5C), in line with patterns in the observational data (Fig. 2B).

**Fig. 5.**
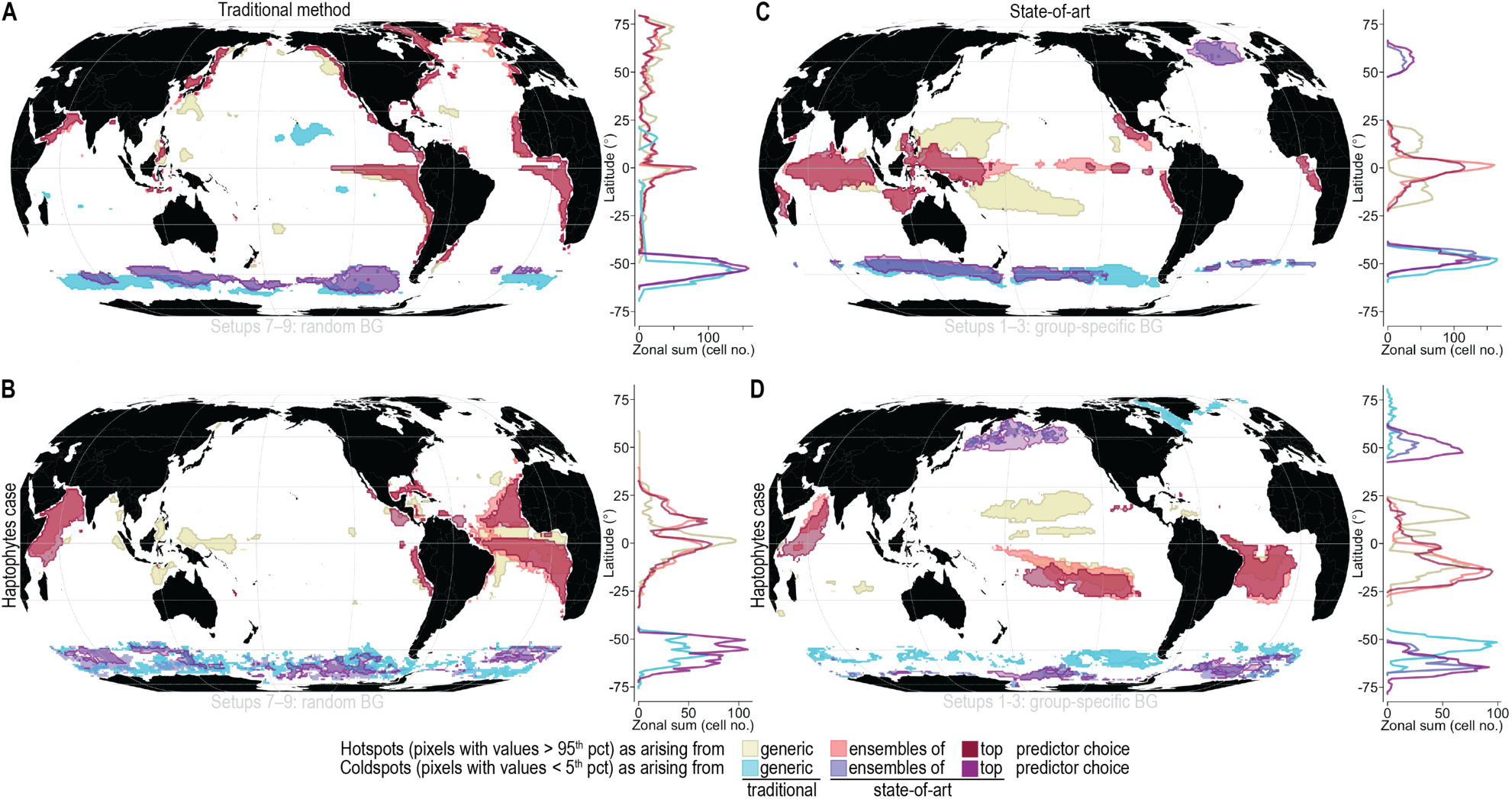
Sensitivity of global species richness hotspots and coldspots to predictor choice using generic versus species-specific selection. Each global map shows hotspots (yellow-red colors) and coldspots (blue colors) arising from alternative predictor choices, for models using traditional (A–B) or advanced (C–D) background data. Results are based on the annual mean maps of monthly phytoplankton species richness of all groups (upper maps) and a particularly data-sparse group (lower maps). Lines next to the maps show the sum of cells per degree latitude that fall into hot/cold-spots per choice.

Across global temperature values, the model setups 1, 4 and 9, which use predictor-ensembles tuned for each species, match the observed ln (richness) slopes better than the generic and top-member choices. The ensemble choice roughly halves observed-to-predicted slope differences compared to the generic choice (fig. S6), holding other factors fixed.

Similarly, as model test scales are widened from monthly to multi-monthly resolution, setups using the ensemble predictor approach retain their predictive skill to a higher degree than setups using the generic predictor set, regardless of the CV metric used (fig. S9, robustness in fig. S10).

#### Projection choice

Presence-absence projection choice leads to significantly higher scores than probabilistic projection choice as assessed by sbCV (Fig. 3B; *p* < 0.001 and *n* = 207 in each algorithm) in all metrics except Pearson’s *r* (see Results in the *SI*). This result is confirmed by the greater ability of PA models than PRB models to reproduce the steepness of latitudinal gradients observed (fig. S3) and the superior capacity of PA models to replicate ln (richness) slopes globally throughout all setups, algorithms and analysis scales (fig. S6).

#### Higher-performing choices

Among the refined background choices, the group-specific (GSTG) choice significantly outperforms the TTG choice (*p* <0.001, all metrics; *n* = 414; 6 setups × 23 scales × 3 algorithms with GSTG vs. TTG) although absolute differences are small (Fig. 3B). Among the two refined predictor choices, the top-member choice receives significantly higher scores than ensemble choice (*p* <0.001 and *n* = 138 for each algorithm). Nevertheless, when assessing global model skill through global richness slopes, setup 1, which uses the full predictor ensembles at species-level, outperforms all other setups. PA-choice, combined with GSTG, and top-member predictor variables yields the SDM setup 2, which receives the overall highest Spearman’s *ρ* mean (SD) score of 0.901 (0.038). In essence, setup 1 and 2 emerge as best performing, depending on the model skill test applied.

## Discussion

Observational data are sparse and lack systematic global design for the vast majority of known species (Mora *et al*. 2011; Meyer *et al*. 2015; Menegotto & Rangel 2018). The modeling choices and multiscale validation method reported here may facilitate the mapping of significant components of under-sampled biodiversity in the global ocean.

Our systematic comparison and cross-validation of model calibration setups offers a baseline for improving biodiversity analyses across global oceans with sparse data that are prone to strong survey bias. Our recommendations to overcome these limitations include (1) the choice of predictors selected from broad sets drivers accounting for gaps in our knowledge of species’ ecology, (2) a careful target-group definition capturing original survey biases for background contrasts, and (3) thresholding of SDMs for global species richness predictions. Choices regarding background data (pseudo-absences) selection influence the outcome of traditional SDM test-scores, such as the AUC or true skill statistic, as these metrics consider absence data in the test. Our results demonstrate the failure of a common random CV approach to assess global SDM skill across a range of background choices in the context of survey bias. We propose a spatial CV approach that circumvents the need for true absence data by using a systematic suite of spatiotemporally extents to analyze diversity patterns in sparse observational data. This avoids perception bias induced by any single scale of analysis (Levin 1992) and allows us to assess global model generalization skill.

Our CV approach addresses further the problem of temporal data sparseness, which aggravates our ability to resolve global biodiversity structure in the context of seasonally highly dynamic microbial organisms. Our data analysis and validation at multiple monthly timescales allowed for successful tests and aggregation of richness patterns. This approach may be transferable to models of other short-lived organisms such as viruses, bacteria, fungi, but also insects, migratory animals, birds, and further biota inactive on sub-annual scales.

The results of our study highlight the capacity of statistical models to uncover patterns, hotspots, and environmental correlates of diversity in the pelagic ocean where biodiversity is often temporally dynamic, yet highly undersampled. We may thus largely overcome the problem of sampling bias (Bardon *et al*. 2021) and data sparseness (Menegotto & Rangel 2018) through advances in model setups. Skilled species distribution models have been determined and used here as a tool that fills observational gaps while paying close attention to data structure. Our results offer a key reference for using niche models with globally sparse data and show how model success depends on partially novel calibration choices that address essential challenges of data-sparse systems. Such refined model setups may enable predictions of global biodiversity patterns for other understudied and under-sampled marine taxa. The accurate mapping of this biodiversity is a key prerequisite to test (macro-)ecological hypotheses, to study functional relationships, infer ecosystem services, and assess conservation needs.

The target-group based selection of background data has been viewed as an important strategy to cope with presence data survey bias (Phillips *et al*. 2009; Ballesteros-Mejia *et al*. 2017), an approach that has recently been used successfully to capture gradients in plankton diversity (Righetti *et al*. 2019; Benedetti *et al*. 2021). However, one uncertain aspect has been the size of the target group. Here, in the presence of global survey bias, we have shown the potential for model skill improvement by refining the target group to the taxon examined. Such refinement may depend on the size of the study group and differences in available sampling coverage among studied taxa. Concise target groups might be useful for the analysis of size structured imaging data where different optical instruments detect specific size ranges or ‘omics information’ targeting varying size classes (Frémont *et al*. 2022; Irisson *et al*. 2022). Importantly, the target group needs to be defined large enough to reflect the environmental conditions of the study area (Guisan & Zimmermann 2000), yet small enough to ascertain that the target group is reflecting the original survey design applied to the model species (Phillips *et al*. 2009). Target-group optimization for background definition exerted massive improvements (3.9 to 21.5-fold compared to random background choice), leading to the most significant model skill gains in our global marine phytoplankton example.

Optimized predictor variable choices allowed for a refined mapping of richness hotspots. Knowledge of skillful predictors to capture the niche of a particular species well is often incomplete, yet defines an important aspect of good SDM practice (Araújo *et al*. 2019). We show that species-specific predictor selection in the presence of unknown elusive niche factors significantly improves model performance over constant predictors carefully selected from key literature (e.g., Brun *et al*. 2015; Barton *et al*. 2016). Uncertainty about the importance of factors shaping ecological niches has been previously identified as a key problem with organisms for which data are sparse (Breiner *et al*. 2015). Our evidence for reduced patchiness in global predictions by SDMs that favor predictor-ensembles over generic predictors suggests that variable sets that address species-level ecological uncertainty may also reduce the impact of survey artefacts and clustered data on predicted richness.

The use of probabilistic species-projections from SDMs has been hypothesized to outperform presence-absence projections when modeling species richness from stacked SDMs (Calabrese *et al*. 2014), and this claim is often replicated and debated (Zurell *et al*. 2016, 2020) also in the context of marine zooplankton (Benedetti *et al*. 2021). We find that stacked presence-absence projections capture latitudinal species richness variations and slopes at a higher accuracy than probability-based projections. One possible reason is that open ocean taxa may be less influenced by local biotic and habitat effects than terrestrial taxa (Guisan & Rahbek 2011). Another reason might be that high latitudes offer cold-water habitats readily captured by SDMs, yielding higher average probabilities of species’ presence in these areas.

The extent to which the lack of explicit representation of biotic interactions in SDMs affects predictions of richness has been debated (Wisz *et al*. 2013). At our scales of analysis, this aspect had a minor effect on prediction accuracy, as our SDMs (not explicitly representing biotic interactions) explained the richness found directly in the observational data (mirroring biotic and abiotic influences) highly faithfully. Yet we cannot exclude the possibility that they may play a role at the level of some individual species.

In summary, on the basis of 1 million open ocean occurrences for 567 phytoplankton species, we offer prospects and guidance on how sparse and biased sampling efforts can be used in a in a way that is applicable to a vast fraction of neglected marine species. The reported concepts may be adapted to species groups that are prone to strong spatiotemporal undersampling and survey bias, providing insights into the structure of global marine biodiversity and offering a wealth of applications from theory testing to conservation.

## Supplementary Information is available for this paper

### Data accessibility statement

Phytoplankton occurrence data underlying this study are available through https://doi.org/10.1594/PANGAEA.904397. Additional data, scripts, and materials related to this paper may be requested from the authors.

## Acknowledgements

We thank D. Loher for aid in cluster computations and M. Lindegren, F. Benedetti, D. Eriksson, Y. Chauvier, A. Psomas and P. Brun for exchange and discussion. We thank the GBIF, OBIS and MAREDAT initiatives and all taxonomists for their efforts to mobilize phytoplankton data. Model designs were supported by ETH Zürich grant ETH-52 13-2 and in part by BNP Paribas grant Notion. Model fitting and cross validation setups were supported by the Swiss National Science Foundation grant P500PB_203241. This project provides an important basis for the goals of the European Union’s Horizon 2020 Research and Innovation Programme under grant no. 862923 (AtlantECO).

## Supplementary Information for

### Box S1. Challenges associated with observational phytoplankton data

The full integration of available data records is highly desirable to achieve skillful species distribution models, especially for species with sparse sampling points. For marine phytoplankton, observational records remain limited over vast regions and a majority of these data are featuring the following notable issues.

#### Taxonomic and spatio-temporal sampling bias

Digital occurrence data that have been mobilized via data-sharing efforts, such as GBIF (www.gbifg.org) and OBIS (www.obis.org) are often taxonomically incomplete, meaning that samples collected from a particular site suffer from species under-detection. Marine phytoplankton species of low abundances are often neglected by non-genetic sampling methodologies (Ser-Giacomi *et al*. 2018), with about 35 to 70% of the local species richness remaining unseen by traditional samples of 5 to 100 mL seawater (Cermeño *et al*. 2014). In addition, samples of phytoplankton are spatially and temporally biased, owing to uneven distributions of survey efforts across the ocean’s surface and seasons (Righetti *et al*. 2020). More generally, marine assemblages in open oceans have been regularly under-sampled in tropical areas and remote regions, where access to field sites is particularly challenging, leading to a “tropical pelagic data gap” (Menegotto & Rangel 2018).

#### Missing absences

Due to the difficulty of confirming a phytoplankton species’ absence and the consequent widespread lack of reliable absences, background data or pseudoabsences are typically required by species distribution models (Barbet-Massin *et al*. 2012). The method designed to sample these background data can be a powerful means to address original sampling bias and address original data gaps, via so-called “target groups” (Phillips *et al*. 2009). This target group denotes the general group or taxon of species studied and is expected to feature a sampling bias that reflects the bias of the species modeled. By sampling data from this target group, the sampling bias underlying the species’ presence data is reduced by introducing a similar bias to the species’ background. The definition of target groups for phytoplankton has been unclear, as phytoplankton span a wide range of phyla and taxonomic groups. Key phyla, such as diatoms and haptophytes, have exhibited differing historical sampling schemes (Righetti *et al*. 2020), which motivates the need to use differing target groups for their species.

#### Temporal species dynamics

Data limitations in phytoplankton are exacerbated by the fact that phytoplankton are generally short lived, and feature often pulsed growth phases or blooms lasting from only days to weeks (Leblanc *et al*. 2018). Therefore, phytoplankton species may appear transiently, especially in seasonal oceans, over the course of the year (Righetti *et al*. 2019). Such transiency is only captured by models that use time discrete matchups between plankton data and environmental conditions (Pinkernell & Beszteri 2014) or predictors capturing temporal variability (Iglesias-Rodríguez *et al*. 2002), while relating phytoplankton observations to annual environmental predictor averages may be inappropriate (Araújo *et al*. 2019).

#### Uncertainty in species’ niche determinants

Species with sparse data are often poorly studied with regards to their physiological tradeoffs and ecological limitations. The selection of skillful predictors for models that capture the ecological niche of a species is therefore typically unclear, yet constitutes an important aspect of good SDM practice (Araújo *et al*. 2019). For rare species, uncertainty about predictor importance has been identified as a key problem (Breiner *et al*. 2015) and predictors in phytoplankton have been debated: while pCO_2_ was proposed as the main driver of calcifying marine phytoplankton (Rivero-Calle *et al*. 2015), another study suggested temperature as the decisive driver (Beaugrand *et al*. 2013). Other possible key drivers include macronutrients (e.g., silicic acid, nitrate, phosphate, iron), water mixing depth, sea surface wind stress, light, and salinity, among others (Irwin *et al*. 2012; Brun *et al*. 2015; Rivero-Calle *et al*. 2015). However, these factors may be of varying importance, depending on the species.

## Supplementary Methods

### Section S1. Spatial data thinning aimed at reducing presence survey bias

We tested data thinning to 150 km and 300 km as a parallel strategy to reduce spatial sampling bias (Aiello-Lammens *et al*. 2015). We discarded this strategy, as many of our, already data-sparse, species retained ≤ 24 presences, which we considered as lower threshold for inclusion in SDMs.

### Section S2. Single-predictor model setups for candidate predictor tests

GAM used smoothing terms with five basis dimensions for their predictor variable. GLM included linear and quadratic terms. To equalize the total weight of presences versus background data per model, background data in GAM and GLM were weighted by the ratio of species’ presence to background points. RF used 300 trees. Weighting of each RF tree was achieved by randomly subsampling same amounts of background points as the species had presence points.

### Section S3. Selection of four generic predictors for SDMs

Based on single-predictor test, we extracted the top-ten ranked (mean across species) predictors as the basis for selection. These predictors included (top to lowest ranked): T, SSW, dMLD1/dt, P, S, pCO_2_, log P, PAR, Si*, N* (see Table S2). We selected monthly sea surface temperature (T), as it represents the top ranked variable, and a likely impactful driver for the distribution of many species (e.g., Beaugrand *et al*. 2013; Righetti *et al*. 2019; Benedetti *et al*. 2021). To represent physical turbulence (Margalef 1978) we included sea surface wind stress, ranked second. We next included pCO_2_ as likely key driver (Rivero-Calle *et al*. 2015), which acts in conjunct with other environmental factors on phytoplankton (Seifert *et al*. 2020). Finally, to represent macronutrient levels, essential for phytoplankton growth, we used N* (Gruber & Sarmiento 1997), as this variable was very weakly correlated with temperature (Table S3). We tested the robustness of the emerging top-ten ranked predictors to background data choice during single-predictor tests. With the exception of a single variable (loss of PAR), the same predictors resulted in the top ten, using TTG rather than GSTG background data.

### Section S4. Scheme to convert globally biased presence data into robust species richness data

To obtain test-data (‘observed truth’) of species richness from globally biased and sparse species’ presence data, we randomly selected and pooled available samples in a spatiotemporally controlled, equal-effort manner, using: (A) constant bandwidths to split the global ocean into latitudinal bands, serving as blocks in the cross-validation, (B) constant amounts of spatiotemporally defined cells selected per band, (C) constant amounts of samples selected per cell, and (D) a taxonomic minimum resolution-threshold for sample acceptance for analyses. This basic design exerts firm control over spatial and temporal sampling effort, while filtering individual samples with particularly low detection ability or unrealistically few species reported.

### Section S5. Specific choices made for test-data generation

We build on the scheme proposed under Section S1 in a comprehensive manner, using 30 scales of sample selection, data aggregation, and data analysis, at high folds (1000 randomized selections). Our choices include: (A) twelve 15° latitude bands spanning the entire globe, (B) fixed amounts of monthly 1°×1° cells selected per 15°-latitude band, at random and without replacement, and (C) fixed amounts of samples drawn from each of the selected monthly 1°×1° cells, at random, and without replacement, and (D) consideration of samples containing ≥ 5 species. We test 30 combinations of cell no. × sample no. in a systematic manner (Fig. 2B).

Regarding (A): Our choice to use latitude bands as the baseline scheme for spatial block cross-validation is motivated by the necessity to bin phytoplankton sample-data into larger units to estimate the species richness (see Cermeño et al., 2014). Latitude bands were preferred over longitude bands, since phytoplankton richness often exhibits distinct latitudinal gradients (Ibarbalz et al. 2019, Righetti et al. 2019b), useful for model testing, at least at sampling volumes larger than conventional volumes. Our choice of 15°-latitude bands reflects a tradeoff between enhancing analysis detail (at smaller bandwidths), while avoiding data gaps (smaller bandwidths would quickly render many bands data-deficient, when aggregating useful amounts of samples).

Regarding (B) and (C): Our choice to use multiple data aggregation scales addresses the scale-sensitivity of observable species richness (Levin 1992; Nogués-Bravo *et al*. 2008). We pooled samples to varying degrees, selecting *m* (ranging from 2 to 10) samples per monthly 1°-cells, times *n* (1 to 6) monthly 1°-cells per latitude band. This matrix covers varying sampling intensity per monthly 1° cells (Fig. 2C; increasing from bottom to top) and varying temporal integration of samples from one, and up to six, monthly cells (Fig. 2C). While an integration of many samples is desirable to improve richness analysis (Cermeño et al. 2014), our possibilities were limited by the lack of samples available, especially in the Southern Hemisphere (Fig. 2B; white scales lacked data). Reaching the upper limits of data availability, we pooled a maximum of 60 samples per band, while keeping resampling effort constant across bands. At even higher efforts (>10 samples per cell, >6 cells per band) many bands were rendered data-deficient.

Regarding (D): Our choice to consider only samples featuring ≥5 species for test data generation, was motivated by a data-detail vs. data-availability tradeoff. We initially explored the richness from pooled samples, relaxing any criterion with respect to the number of species per sample (*n* samples = 182’303). This strategy led to strong noise and erratically low tropical species richness estimates (see Fig. S2). We therefore introduced a minimum threshold. Our final choice of ≥5 species per sample (*n* samples = 63’441) represents a tradeoff between avoiding low taxonomic information contained per sample (that is, excluding samples with only one species, or samples with up to four species; compare with e.g. Cermeño et al. 2014, Rodríguez-Ramos et al. 2015, Ibarbalz et al. 2019), while retaining sufficient samples for global test data generation. The choice of ≥5 species per sample led to an omission of 65.2% of original samples, and 27.4% of original presences. Higher thresholds (e.g., ≥6 species or ≥7 species per sample) led to the omission of substantial data (72.0% or 77.32% of original samples, respectively). Unlike test data, our SDMs considered the full data (samples with any species numbers) for the purpose of global species richness modeling, being inclusive of the full empirical samples (of any quality) available.

### Section S6. Spatiotemporal matchup of predicted with observational data-derived richness

A key aspect of our spatial block CV is the tight comparison of species richness predicted to that observed in test-data, unique to each of the 1000 selection folds, per 23 spatiotemporal analysis scales, and 18 SDM setups. Specifically, for each of the 1000 selection folds, and each 15° band, we noted the identities of the monthly 1° cells, selected at random to generate the test-data. When pooling the predicted richness across cells selected per 15° band, the exact same cells, as used for estimating richness in pooled observations, were considered. That is, predictions were tailored on the 1° cells contributing test-data. By computing predicted richness for 1000 selection folds, and then drawing the average, the modeled trend matched the ‘observed truth’.

While the 15° latitude bands represent a rather wide unit to estimate species richness, we nevertheless achieve high spatiotemporal detail in predicted-to-observed contrasts (both latitudinally and longitudinally), as the explicit nature of our comparison that builds on a few monthly 1°-cells, drawn from 15°-bands, per fold.

### Section S7. CV metrics and the standardization of richness values

To measure similarity between observed and predicted richness, we employed commonly used regression and correlation based metrics. The mean Absolute Error (MAE) represents a regression-based error metric, quantifying the mean absolute residuals of a linear model (i.e., ordinary least squared linear-model fit to predicted versus observed richness). In our case, we forced the fit through the origin, arguing that zero modeled and zero observational richness are coincident. By contrast, Spearman’s *ρ*, and Pearson’s *r* describe the correlations between observational and modeled richness, without invoking any regression fits.

To calculate the Spearman’s *ρ*, and Pearson’s *r*, the richness values from models and observations were used straightly, while for MAE, RMSE, we standardized the maxima of observed and modeled latitudinal richness to one. This allowed for comparisons of CV scores across the 23 analysis scales, between which absolute richness values differed, given the varying sample integration efforts between scales (Fig. 2).

## Supplementary Results

We elaborated statistically superior SDM choices with regards to background selection and predictor variable choice further.

### Superior choices of background data

Target-group background choices reach substantially higher CV scores than the random background choice (Fig. 3B; paired Wilcoxon test, *p* <0.001, all metrics). Among the two target group choices, the GSTG choice significantly outperforms the TTG choice (*p* <0.001, all metrics; *n* = 414; 6 setups × 23 scales × 3 algorithms with GSTG vs TTG) although absolute differences are minor (Fig. 3B; compare gray shades). Analyzed by algorithm, the difference remains highly significant for RF (*p* <0.001, all metrics, *n* = 138), and partially highly significant for GAM (*ρ* ns, other metrics *p* <0.001, *n* = 138) and GLM-fitted SDMs (*p* ≤0.001 in *r* and *ρ*; ns in MAE; *p* = 0.18 in RMSE, *n* = 138). Accordingly, prediction biases relative to the “observed truth” are generally low for target-group backgrounds (Fig. 3A; setups 1–6) but strongly amplified for the random background (Fig. 3A; setups 7–9), especially >30°N or at 40°S, which is reflected in poor SDM skill (Fig. 3B).

### Superior choices of predictor variables

Based on the *ρ* metric, our species-specific predictor choices improve SDMs significantly relative to the generic predictor choice (*p* <0.001 and *n*=138 for each algorithm except RF, ns). Among the two species-specific choices, the top member choice reaches significantly higher scores than top ensemble choice (*p* <0.001 and *n* = 138, for each algorithm). Specific-specific choices improve the scores particularly clearly for GAM and GLM when combined with the random “baseline” background choice (Fig. 3B, Table S3).

For the remaining CV metrics, results are mixed. While MAE and RMSE scores are enhanced by species-specific predictor choices in GAM (p <0.08 or p <0.22, respectively, n = 138) and partially enhanced in GLM (top member p <0.06, else ns; n = 138), they are worsened in RF (*p* <0.001, *n*=138). For the *r* metric, species-specific predictor choices lead to improved scores relative to generic ones in GAM (*p* <0.001, *n* = 138) but there are no significant differences in GLM and lower scores in RF (*p* <0.001, *n* = 138).

**Fig. S1.**
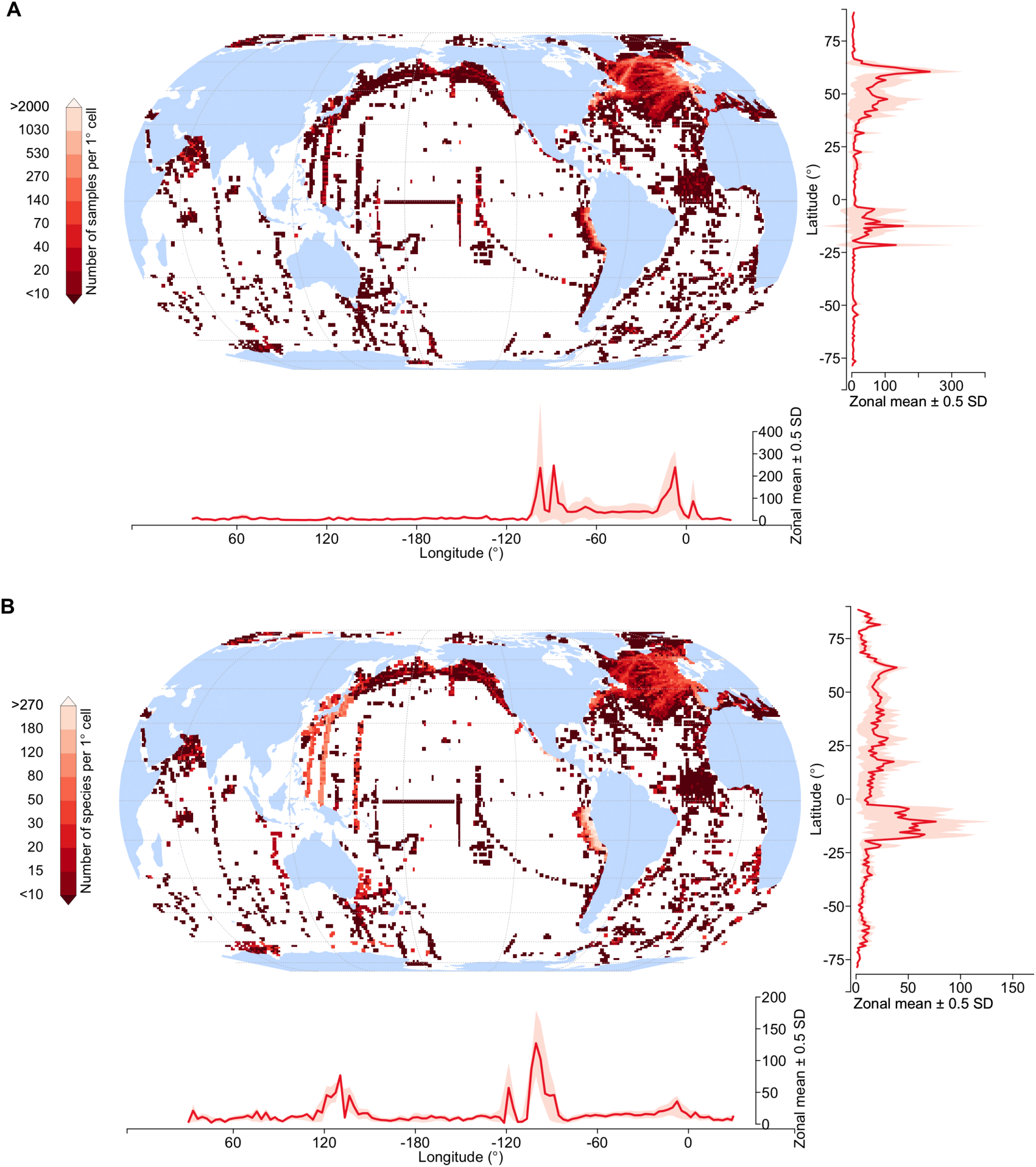
Global distribution of sampling effort and species richness detected per 1° cells. (A) Number of phytoplankton in situ samples available per 1° × 1° grid cell, reflecting cumulative sampling efforts, and the species richness (B) detected per 1° × 1° grid cell in consequence of these efforts (Pearson’s *r* of A vs B = 0.44). Gradients for maps show the means (0.5 SD in shade) per degree latitude or longitude, omitting cells lacking data. The monthly open ocean 1° cells containing samples (*n* = 17’333) cover just 3.5% of all monthly open ocean 1° cells, and 42.1% of open ocean 1°cells when discarding monthly distinction. Data binned per monthly 1° cells provided the basis for generating the global (latitude-longitude-month) diversity predictions presented in this study. Refer to Figure 2A for spatial coverage of samples used for richness test data generation.

**Fig. S2.**
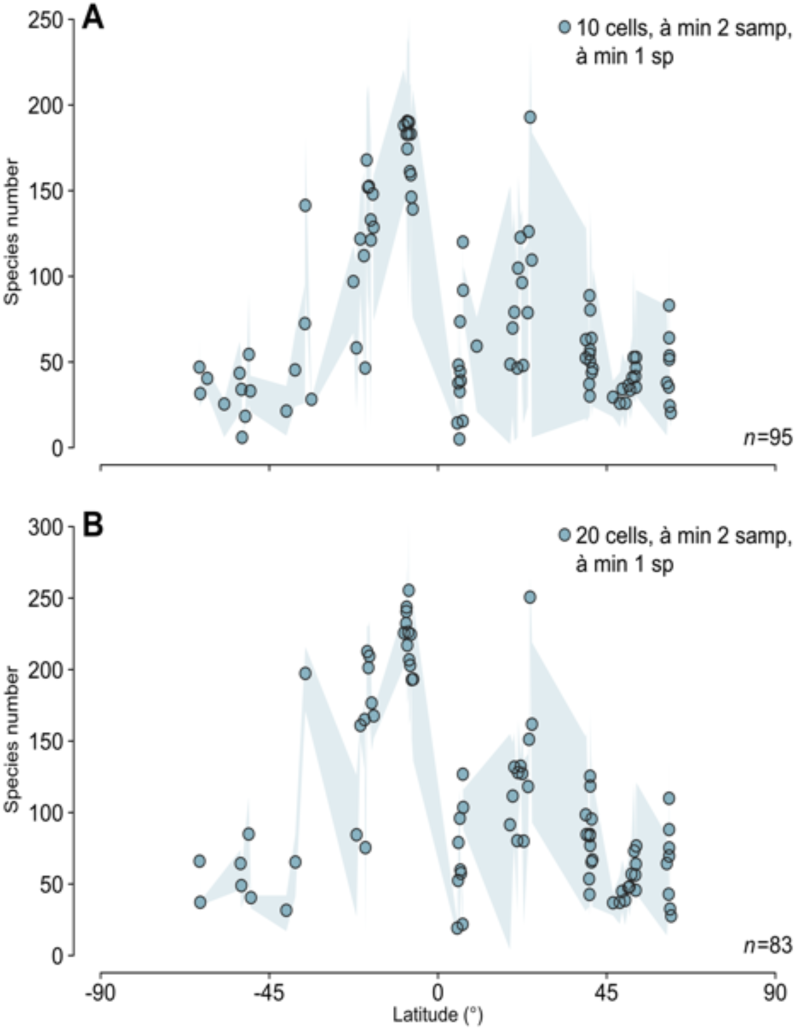
Precursor method used to extract species richness gradients in the observations. Dots show the mean of species richness obtained from data pooled per monthly 15° latitude band and calendarial month, and shadings the 5^th^ to 95^th^ percentile ranges of the 1000 folds. In panel (A), ten 1° cells, and in panel (B), twenty 1° cells were selected at random (from the cells containing ≥ 2 samples) per 15° latitude monthly band (spatial band definition, see Fig. 2A). In contrast to the final method used (see Figure 2), here, all samples per 1° cell selected were retained and no lower threshold regarding species number is applied to samples. Such failure to correct for equal amounts of samples selected per cell, and the inclusion of samples with low (2–4) species resulted in rather erratic latitudinal diversity estimates with inflated tropical richness at ∼10° North.

**Fig. S3.**
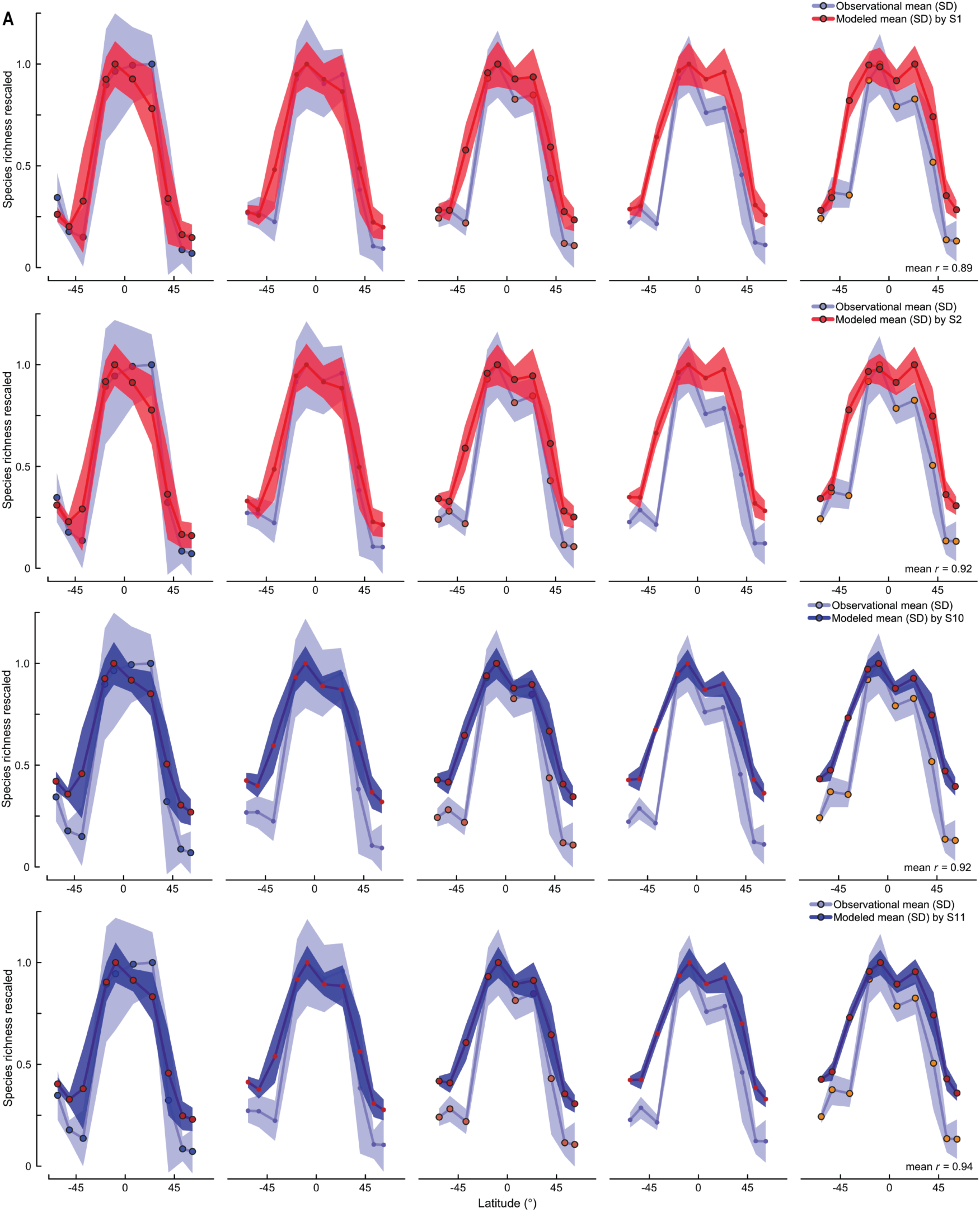
Predicted-to-observed richness contrasts for the top ranked models (RF) by Spearman’s *ρ*score, showing PA models in red (Setup1 and 2) vs. PRB models in blue (Setup10 and 11). Data show richness at five exemplary spatiotemporal scales (see Fig. 2B). Although PRB models show strongest correlation with observations, they feature short “legs” (i.e., shallower latitudinal gradients) relative to PA models and observation.

**Fig. S4.**
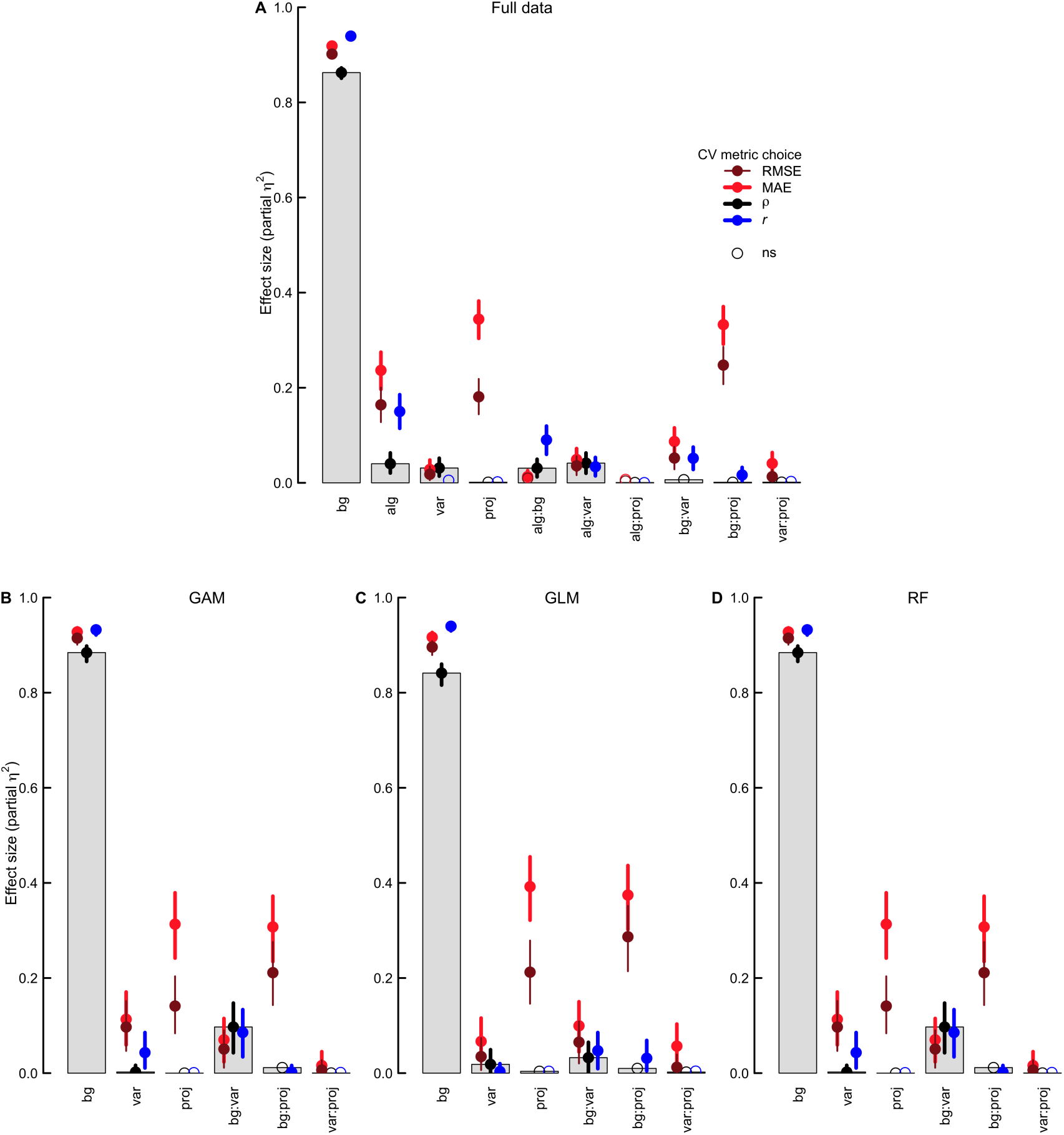
Effect size of methodological choices on global SDM skill. Effect sizes are estimated by ANOVA for four individual cross validation (CV) metrics. (A) Results emerging from the full data (18 SDM setups × 23 analysis scales x 3 algorithms). (B–C) Results within individual algorithms. The CV metrics assessed the skill of SDM setups via predicted-to-observed species richness contrasts, using spatial-block CV. Filled dots represent the magnitude of effects on SDM predictive skill for significant effects (*p* <0.001). Vertical lines denote 95% confidence intervals. Model calibration choices included: bg, background; alg, fitting algorithm; var, predictor variables; proj, projection type, and their linear combinations. Residual effects were negligible.

**Fig. S5.**
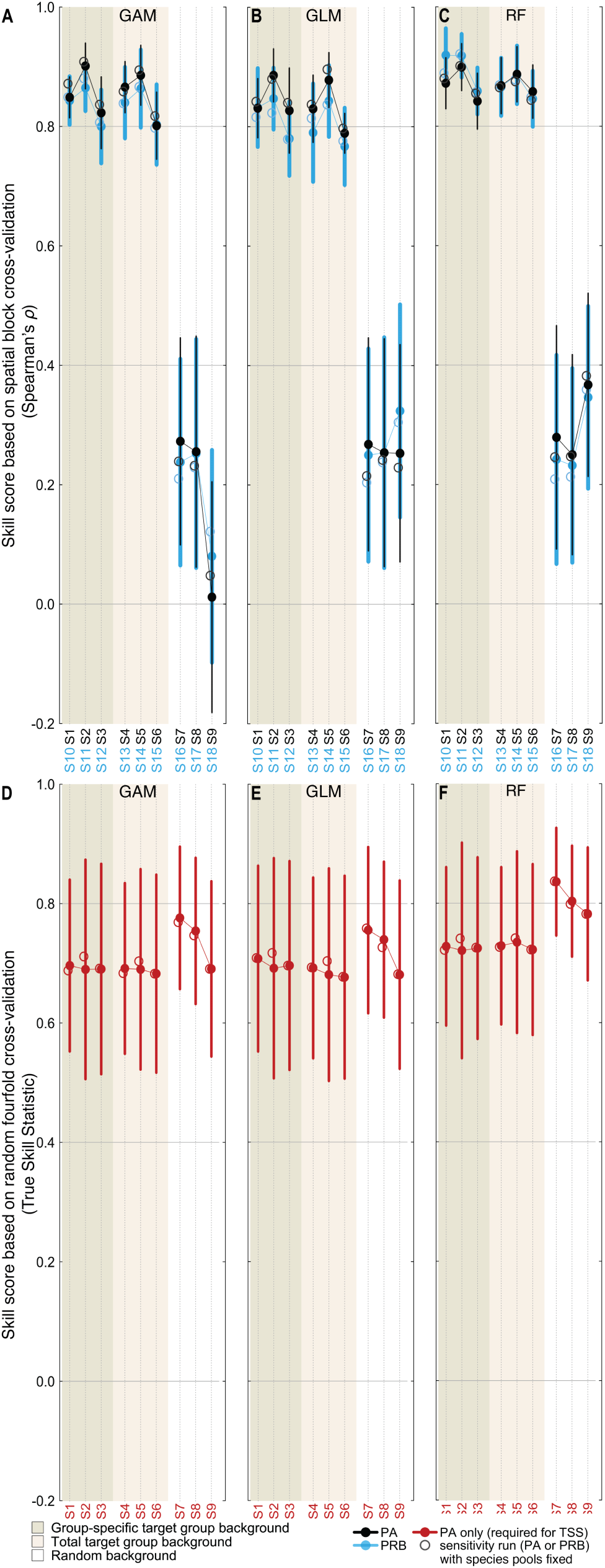
SDM predictive skill as assessed by multiscale spatial block CV and random fourfold CV for each algorithm tested in this study. Filled circles indicate results based on all successfully modeled species. Open circles show the results of a sensitivity run, correcting for equal species pools in richness estimation and evaluation. In panels (A– C), dots are means (lines for SD) of 23 analysis scales that test SDM performance globally at richness level, while in panels (D–F), dots are means (lines for SD) of species’ TSS scores achieved by SDMs (single SDMs for S2, 3, 5, 6 and 8, 9, and corresponding PRB setups in light blue font, and direct evaluation of ensembles of SDMs for S1, 4, and 7, and corresponding PRB setups in light blue font). Members of ensembles have been binarized to 0/1 prior to member-averaging during fourfold CV, using thresholds maximizing TSS, all else equal to four-fold CV for single SDMs (Methods). Note that TSS scores require a thresholding of models to 0/1 and thus can be determined for PA model setups only.

**Fig. S6.**
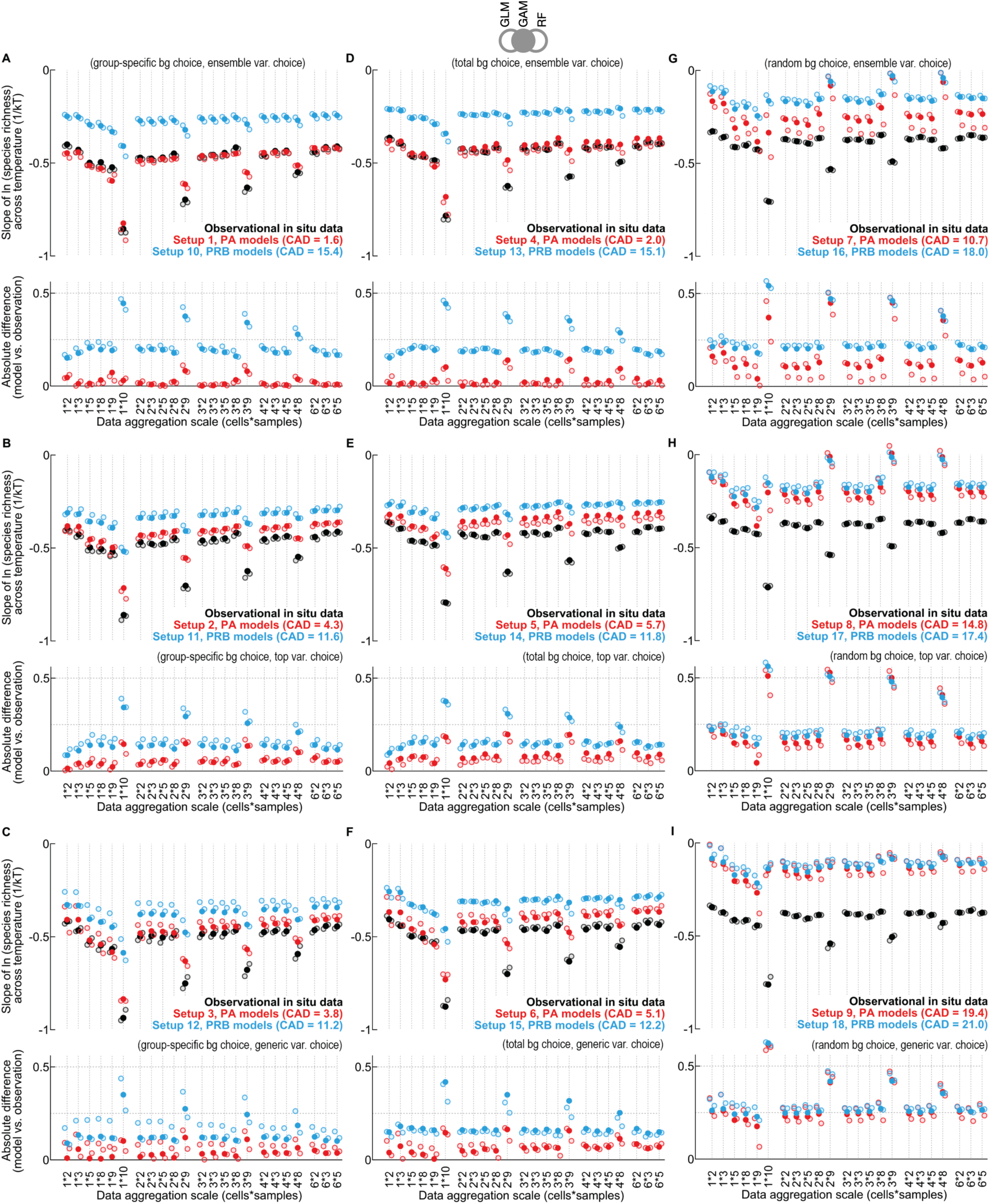
Model performance in predicting linear slopes of richness across temperature. The natural logarithm (ln) of species richness is contrasted to Temperature (k, Boltzmann’s constant; T, sea surface temperature in Kelvin) for all scales, model setups, and algorithms tested. Dots represent linear slope coefficient fitted on the global species richness across temperature for a particular data aggregation scale (filled dots for GAM, open dots for GLM and RF). Absolute predicted-do-observed difference for a particular scale are inserted at the bottom of each panel/model setup. CAD is the cumulative absolute difference between predicted slopes (colored dots) vs. observed slopes (black). Observed richness data are identical to those used for block CV (Fig. 2). Panels are sorted by background choice (left to right), and variable choice (top to bottom). Sea surface temperature values were matched with corresponding observational data at monthly climatological 1° grid-cell resolution for this analysis, using an averaging per band latitude in line with the pooling of observations used to determine richness, and stem from the World Ocean Atlas 2013 (Locarnini et al., 2013).

**Fig. S7.**
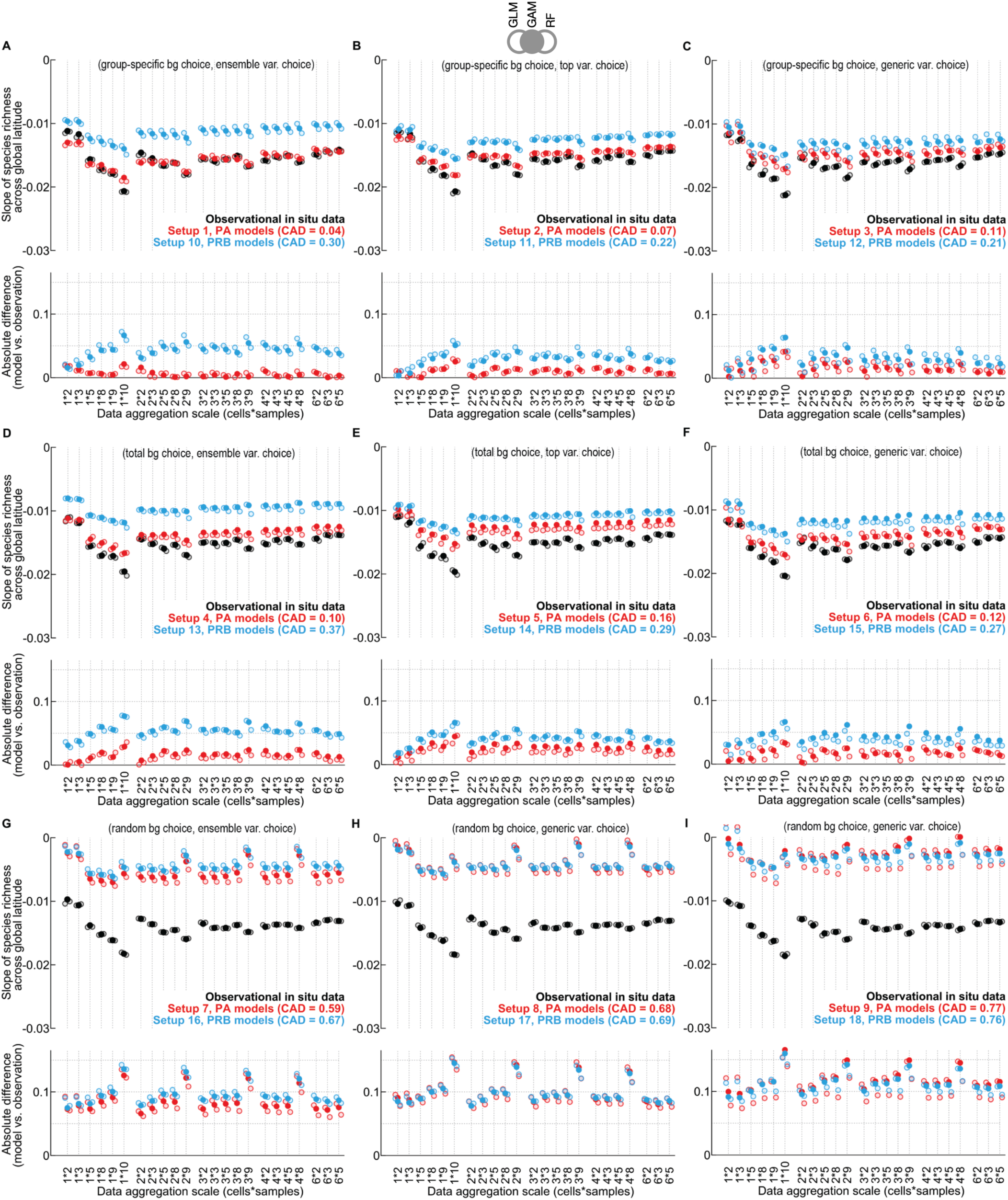
Model performance in predicting linear slopes of richness across global latitude. Species richness is contrasted to absolute latitude (degrees) for all scales, model setups, and algorithms tested. Dots represent linear slope coefficient fitted on the global species richness across latitude for a particular data aggregation scale (filled dots for GAM, open dots for GLM and RF). Absolute predicted-do-observed difference for a particular scale are inserted at the bottom of each panel/model setup. CAD is the cumulative absolute difference between predicted slopes (colored dots) vs. observed slopes (black). Observed richness data are identical to those used for block CV (Fig. 2). Panels are sorted by background choice (left to right), and variable choice (top to bottom). Latitude values are matched with corresponding observational data at 1° resolution for this analysis, using the means per band latitude in line with the pooling of cells containing sampling data used to determine richness.

**Fig. S8.**
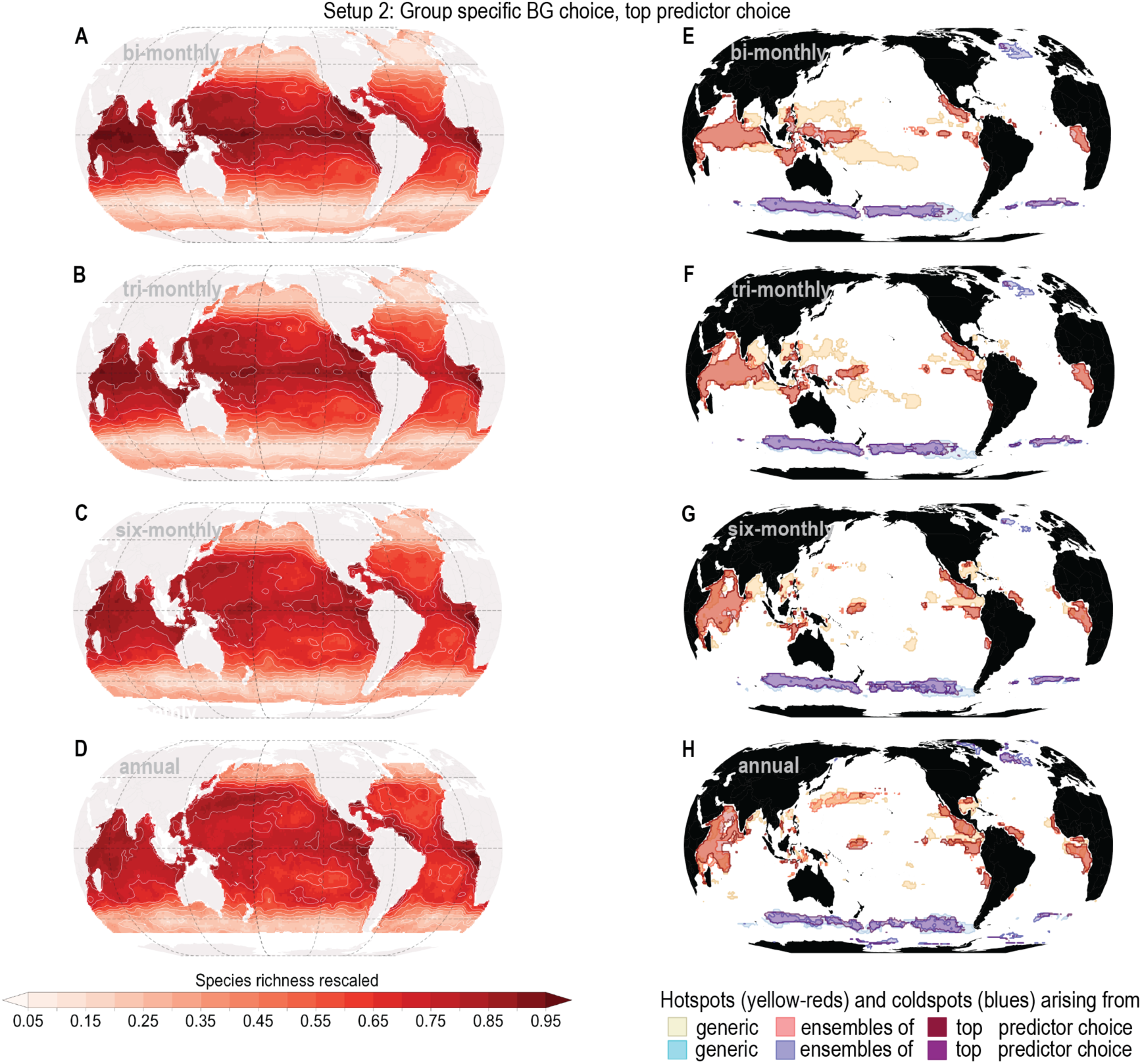
Global patterns and hotspots at varying temporal scales of data integration. (A–D) shows the species richness analyzed at bi-monthly to annual time resolutions. (E–G) shows predicted global hotspots (cells with values above the 95^th^ percentile of richness values) and coldspots (cells with values below the 5^th^ percentile of richness values) under different variable choices explored at bi-monthly to annual time resolutions. The maps show the mean values of the phytoplankton species richness patterns at the defined time-resolutions across the annual cycle (Methods).

**Fig. S9.**
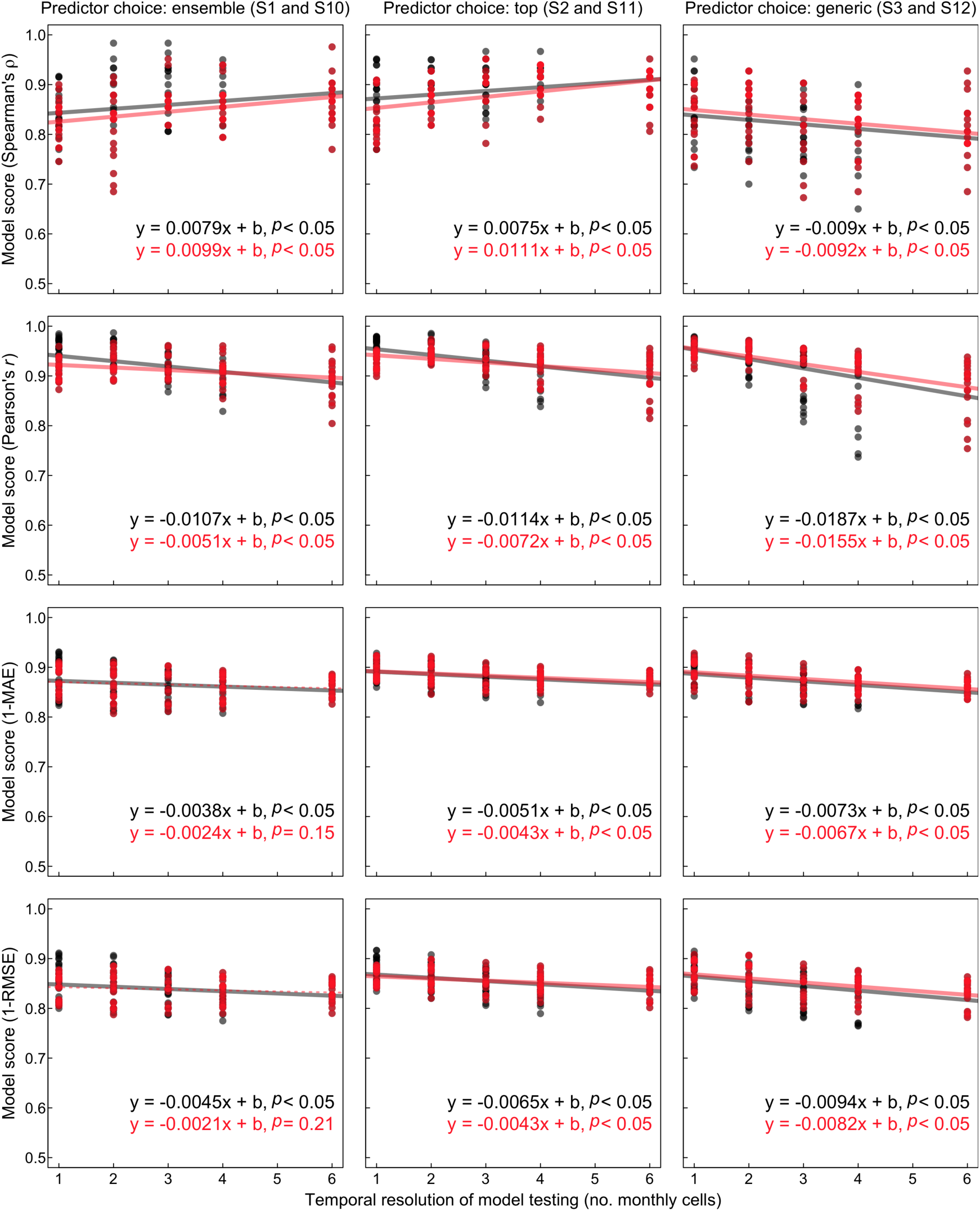
Model skill as a function of the temporal scale of testing. Data are shown for the highly refined target group approach (GSTG). Panels from left to right represent three different variable selection strategies. From top to bottom, different CV metrics are examined. Lines are fitted ordinary least-squared regressions on the full data (black; using all 23 scales of analysis), and for data keeping the resampling effort per cell constant across the five temporal scales of model testing (red; using five scales of analysis, i.e., 5 samples x *m* cells, with *m* equal to 1, 2, 3, 4, and 6; see Figure 2B). The generic predictor choice (right column) leads to estimated slopes on model skill that are consistently more negative than those of the ensemble or top variable choice

**Fig. S10.**
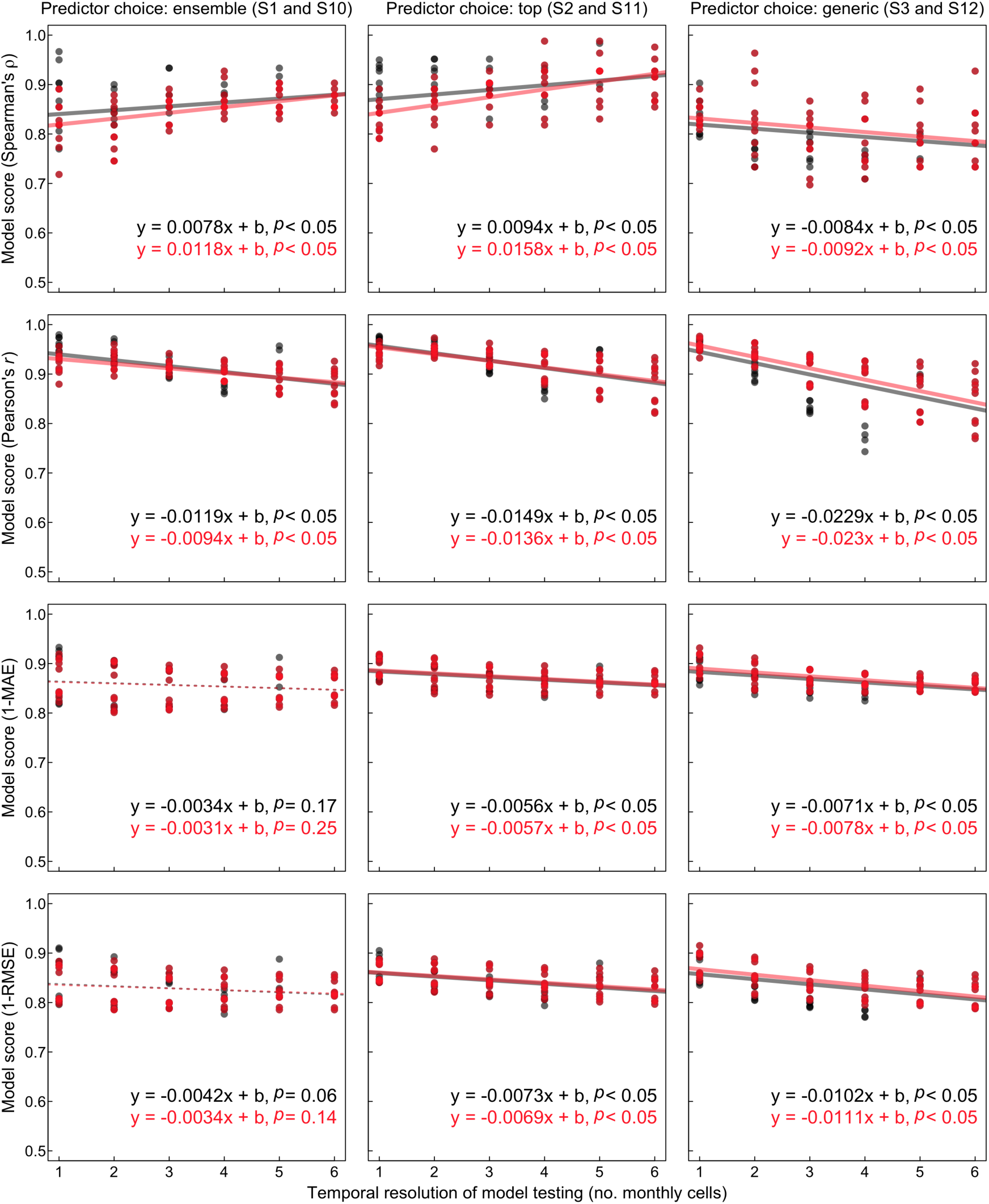
Model skill as a function of the temporal scale of testing under expanded data. Data are shown for the highly refined target group approach (GSTG), as for figure S7. However, to test the robustness of results we used here identical species pools in all model setups (S1–S3, plus S10–12) and a larger suite of 56 test scales (i.e., all scales being filled in the resampling matrix of Figure 2B).

**Table S1.**
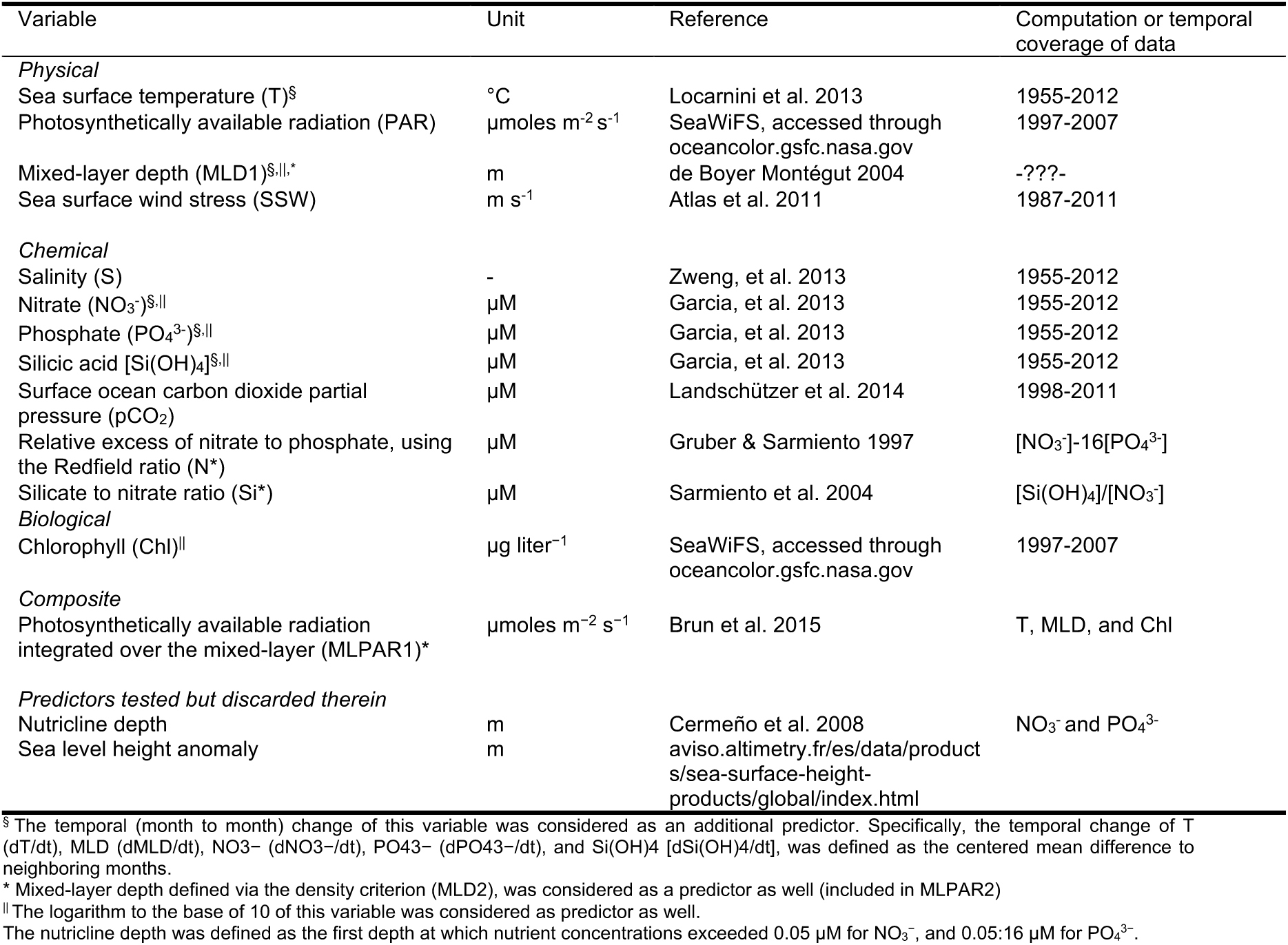
SDM candidate predictor variables.

**Table S2.** Spearman correlations (*ρ*) between predictor variables.

|  | T | S | NO<sub>3</sub><sup>-</sup> | PO<sub>4</sub><sup>3-</sup> | Si(OH)<sub>4</sub> | MLD1 | MLD2 | PAR | Chl | SSW | pCO<sub>2</sub> | MLPAR1 | MLPAR2 | N-star | Si-star | dT/dt | dNO<sub>3</sub><sup>-</sup>/dt | dPO<sub>4</sub><sup>3-</sup>/dt | dSi(OH)<sub>4</sub>/dt | dMLD1/dt | logMLD1 | logMLD2 | log(Chl) | log(NO<sub>3</sub><sup>-</sup>) | log(PO<sub>4</sub><sup>3-</sup>) | log(Si(OH)<sub>4</sub>) |
| --- | --- | --- | --- | --- | --- | --- | --- | --- | --- | --- | --- | --- | --- | --- | --- | --- | --- | --- | --- | --- | --- | --- | --- | --- | --- | --- |
| T | 1 | 0.43 | 0.71 | 0.70 | 0.47 | 0.37 | 0.53 | 0.62 | 0.50 | 0.69 | 0.37 | 0.65 | 0.71 | 0.17 | 0.25 | 0.03 | 0.03 | 0.05 | 0.05 | 0.01 | 0.37 | 0.53 | 0.50 | 0.71 | 0.70 | 0.47 |
| S |  | 1 | 0.52 | 0.57 | 0.53 | 0.07 | 0.11 | 0.39 | 0.37 | 0.27 | 0.07 | 0.32 | 0.33 | 0.23 | 0.02 | 0.02 | 0.01 | 0.03 | 0.04 | 0.03 | 0.07 | 0.11 | 0.37 | 0.52 | 0.57 | 0.53 |
| NO<sub>3</sub><sup>-</sup> |  |  | 1 | 0.82 | 0.57 | 0.34 | 0.43 | 0.48 | 0.57 | 0.56 | 0.02 | 0.59 | 0.61 | 0.19 | 0.37 | 0.02 | 0.02 | 0.06 | 0.05 | 0.00 | 0.34 | 0.43 | 0.57 | 1.00 | 0.82 | 0.57 |
| PO<sub>4</sub><sup>3-</sup> |  |  |  | 1 | 0.57 | 0.35 | 0.43 | 0.45 | 0.53 | 0.51 | 0.04 | 0.56 | 0.58 | 0.57 | 0.29 | 0.05 | 0.02 | 0.05 | 0.05 | 0.02 | 0.35 | 0.43 | 0.53 | 0.82 | 1.00 | 0.57 |
| Si(OH)<sub>4</sub> |  |  |  |  | 1 | 0.16 | 0.22 | 0.34 | 0.43 | 0.33 | 0.01 | 0.37 | 0.39 | 0.18 | 0.31 | 0.04 | 0.03 | 0.05 | 0.05 | 0.04 | 0.16 | 0.22 | 0.43 | 0.57 | 0.57 | 1.00 |
| MLD1 |  |  |  |  |  | 1 | 0.93 | 0.63 | 0.05 | 0.71 | 0.15 | 0.81 | 0.77 | 0.11 | 0.28 | 0.44 | 0.13 | 0.10 | 0.10 | 0.07 | 1.00 | 0.93 | 0.05 | 0.34 | 0.35 | 0.16 |
| MLD2 |  |  |  |  |  |  | 1 | 0.68 | 0.13 | 0.79 | 0.21 | 0.83 | 0.86 | 0.11 | 0.31 | 0.40 | 0.11 | 0.07 | 0.07 | 0.08 | 0.93 | 1.00 | 0.13 | 0.43 | 0.43 | 0.22 |
| PAR |  |  |  |  |  |  |  | 1 | 0.36 | 0.65 | 0.37 | 0.89 | 0.89 | 0.09 | 0.15 | 0.61 | 0.16 | 0.14 | 0.10 | 0.36 | 0.63 | 0.68 | 0.36 | 0.48 | 0.45 | 0.34 |
| Chl |  |  |  |  |  |  |  |  | 1 | 0.31 | 0.20 | 0.50 | 0.51 | 0.24 | 0.09 | 0.03 | 0.06 | 0.05 | 0.06 | 0.09 | 0.05 | 0.13 | 1.00 | 0.57 | 0.53 | 0.43 |
| SSW |  |  |  |  |  |  |  |  |  | 1 | 0.34 | 0.78 | 0.80 | 0.08 | 0.31 | 0.29 | 0.10 | 0.08 | 0.06 | 0.12 | 0.71 | 0.79 | 0.31 | 0.56 | 0.51 | 0.33 |
| pCO<sub>2</sub> |  |  |  |  |  |  |  |  |  |  | 1 | 0.30 | 0.33 | 0.10 | 0.05 | 0.07 | 0.01 | 0.01 | 0.02 | 0.03 | 0.15 | 0.21 | 0.20 | 0.02 | 0.04 | 0.01 |
| MLPAR1 |  |  |  |  |  |  |  |  |  |  |  | 1 | 0.98 | 0.17 | 0.26 | 0.50 | 0.12 | 0.10 | 0.08 | 0.19 | 0.81 | 0.83 | 0.50 | 0.59 | 0.56 | 0.37 |
| MLPAR2 |  |  |  |  |  |  |  |  |  |  |  |  | 1 | 0.17 | 0.27 | 0.47 | 0.11 | 0.08 | 0.06 | 0.18 | 0.77 | 0.86 | 0.51 | 0.61 | 0.58 | 0.39 |
| N-star |  |  |  |  |  |  |  |  |  |  |  |  |  | 1 | 0.02 | 0.01 | 0.02 | 0.01 | 0.02 | 0.05 | 0.11 | 0.11 | 0.24 | 0.19 | 0.57 | 0.18 |
| Si-star |  |  |  |  |  |  |  |  |  |  |  |  |  |  | 1 | 0.01 | 0.00 | 0.02 | 0.01 | 0.03 | 0.28 | 0.31 | 0.09 | 0.37 | 0.29 | 0.31 |
| dT/dt |  |  |  |  |  |  |  |  |  |  |  |  |  |  |  | 1 | 0.32 | 0.31 | 0.21 | 0.62 | 0.44 | 0.40 | 0.03 | 0.02 | 0.05 | 0.04 |
| dNO<sub>3</sub><sup>-</sup>/dt |  |  |  |  |  |  |  |  |  |  |  |  |  |  |  |  | 1 | 0.34 | 0.22 | 0.27 | 0.13 | 0.11 | 0.06 | 0.02 | 0.02 | 0.03 |
| dPO<sub>4</sub><sup>3-</sup>/dt |  |  |  |  |  |  |  |  |  |  |  |  |  |  |  |  |  | 1 | 0.21 | 0.27 | 0.10 | 0.07 | 0.05 | 0.06 | 0.05 | 0.05 |
| dSi(OH)<sub>4</sub>/dt |  |  |  |  |  |  |  |  |  |  |  |  |  |  |  |  |  |  | 1 | 0.17 | 0.10 | 0.07 | 0.06 | 0.05 | 0.05 | 0.05 |
| dMLD1/dt |  |  |  |  |  |  |  |  |  |  |  |  |  |  |  |  |  |  |  | 1 | 0.07 | 0.08 | 0.09 | 0.00 | 0.02 | 0.04 |
| logMLD1 |  |  |  |  |  |  |  |  |  |  |  |  |  |  |  |  |  |  |  |  | 1 | 0.93 | 0.05 | 0.34 | 0.35 | 0.16 |
| logMLD2 |  |  |  |  |  |  |  |  |  |  |  |  |  |  |  |  |  |  |  |  |  | 1 | 0.13 | 0.43 | 0.43 | 0.22 |
| log(Chl) |  |  |  |  |  |  |  |  |  |  |  |  |  |  |  |  |  |  |  |  |  |  | 1 | 0.57 | 0.53 | 0.43 |
| log(NO<sub>3</sub><sup>-</sup>) |  |  |  |  |  |  |  |  |  |  |  |  |  |  |  |  |  |  |  |  |  |  |  | 1 | 0.82 | 0.57 |
| log(PO<sub>4</sub><sup>3-</sup>) |  |  |  |  |  |  |  |  |  |  |  |  |  |  |  |  |  |  |  |  |  |  |  |  | 1 | 0.57 |
| log(Si(OH)<sub>4</sub>) |  |  |  |  |  |  |  |  |  |  |  |  |  |  |  |  |  |  |  |  |  |  |  |  |  | 1 |

Values exceeding |0.7|, considered as maximum threshold within SDMs, are highlighted by red shade. Pairwise correlations are computed on the basis of the monthly climatological 1° × 1° resolution of predictor data across the global open ocean (open ocean definition, see Methods). MLD1 and MLPAR1 use the temperature criterion of mixed-layer depth (de Boyer Montégut 2004). MLD2, and derivatives, use the density criterion. log, logarithm to base 10.

**Table S3.** Summary statistics of the global skill of the 18 SDM setups in reproducing (res, resubstitution) and predicting (pred, block cross-validation) the richness variations (N = 23 gradients) found in pooled observations, using three metrics for each of three algorithms. Each number presents the mean (standard deviation) of the 23 gradients. Higher predictive/reproductive skill is indicated by higher correlation coefficients and lower MAE.

| Setup | GLM *res* | GLM *pred* | %loss | GAM *res* | GAM *pred* | %loss | RF *res* | RF *pred* | %loss |
| --- | --- | --- | --- | --- | --- | --- | --- | --- | --- |
| **Spearman's correlation coefficient ($\rho$)** |  |  |  |  |  |  |  |  |  |
| S1 | 0.850 (0.050) | 0.831 (0.050) | 2.2 | 0.872 (0.047) | 0.849 (0.034) | 2.7 | 0.905 (0.047) | 0.872 (0.043) | 3.6 |
| S2 | 0.888 (0.053) | 0.886 (0.044) | 0.2 | 0.911 (0.038) | 0.901 (0.038) | 1.1 | 0.923 (0.045) | 0.899 (0.039) | 2.6 |
| S3 | 0.747 (0.116) | 0.827 (0.071) | -10.7 | 0.822 (0.086) | 0.823 (0.060) | -0.1 | 0.813 (0.063) | 0.842 (0.047) | -3.6 |
| S4 | 0.838 (0.056) | 0.830 (0.056) | 1.0 | 0.876 (0.046) | 0.866 (0.866) | 1.2 | 0.905 (0.050) | 0.868 (0.047) | 4.1 |
| S5 | 0.898 (0.050) | 0.878 (0.046) | 2.2 | 0.921 (0.050) | 0.886 (0.050) | 3.8 | 0.922 (0.042) | 0.887 (0.046) | 3.8 |
| S6 | 0.800 (0.054) | 0.789 (0.033) | 1.4 | 0.816 (0.039) | 0.801 (0.056) | 1.8 | 0.816 (0.059) | 0.858 (0.045) | -5.1 |
| S7 | 0.326 (0.181) | 0.268 (0.178) | 17.8 | 0.357 (0.157) | 0.273 (0.173) | 23.6 | 0.446 (0.160) | 0.279 (0.187) | 37.4 |
| S8 | 0.330 (0.182) | 0.254 (0.191) | 23 | 0.344 (0.165) | 0.255 (0.193) | 25.8 | 0.387 (0.164) | 0.250 (0.168) | 35.4 |
| S9 | 0.380 (0.170) | 0.253 (0.182) | 33.4 | 0.319 (0.165) | 0.012 (0.193) | 96.3 | 0.455 (0.172) | 0.367 (0.153) | 19.3 |
| S10 | 0.832 (0.067) | 0.832 (0.066) | 0.0 | 0.856 (0.046) | 0.843 (0.040) | 1.5 | 0.920 (0.045) | 0.919 (0.045) | 0.1 |
| S11 | 0.851 (0.051) | 0.847 (0.052) | 0.5 | 0.900 (0.047) | 0.865 (0.039) | 3.9 | 0.934 (0.038) | 0.918 (0.036) | 1.7 |
| S12 | 0.782 (0.063) | 0.780 (0.062) | 0.3 | 0.780 (0.073) | 0.800 (0.062) | -2.6 | 0.897 (0.030) | 0.859 (0.039) | 4.2 |
| S13 | 0.791 (0.080) | 0.790 (0.082) | 0.1 | 0.828 (0.061) | 0.840 (0.060) | -1.4 | 0.894 (0.041) | 0.866 (0.048) | 3.1 |
| S14 | 0.824 (0.070) | 0.843 (0.060) | -2.3 | 0.862 (0.052) | 0.863 (0.065) | -0.2 | 0.918 (0.041) | 0.886 (0.049) | 3.5 |
| S15 | 0.798 (0.069) | 0.767 (0.064) | 3.9 | 0.795 (0.063) | 0.803 (0.067) | -1.0 | 0.859 (0.032) | 0.846 (0.047) | 1.5 |
| S16 | 0.314 (0.165) | 0.250 (0.178) | 20.4 | 0.320 (0.162) | 0.238 (0.173) | 25.7 | 0.456 (0.163) | 0.242 (0.175) | 46.9 |
| S17 | 0.324 (0.176) | 0.254 (0.193) | 21.6 | 0.328 (0.185) | 0.252 (0.192) | 23.1 | 0.403 (0.176) | 0.232 (0.163) | 42.4 |
| S18 | 0.406 (0.172) | 0.324 (0.178) | 20.2 | 0.336 (0.157) | 0.080 (0.178) | 76.2 | 0.481 (0.153) | 0.346 (0.153) | 28.1 |
| **Pearson's correlation coefficient ($r$)** |  |  |  |  |  |  |  |  |  |
| S1 | 0.910 (0.043) | 0.899 (0.041) | 1.2 | 0.927 (0.038) | 0.917 (0.035) | 1.1 | 0.946 (0.027) | 0.931 (0.027) | 1.6 |
| S2 | 0.925 (0.046) | 0.922 (0.040) | 0.3 | 0.938 (0.042) | 0.930 (0.037) | 0.9 | 0.955 (0.027) | 0.949 (0.025) | 0.6 |
| S3 | 0.880 (0.092) | 0.904 (0.069) | -2.7 | 0.907 (0.072) | 0.906 (0.064) | 0.1 | 0.941 (0.035) | 0.939 (0.029) | 0.2 |
| S4 | 0.899 (0.035) | 0.899 (0.036) | 0.0 | 0.915 (0.034) | 0.910 (0.034) | 0.5 | 0.945 (0.028) | 0.914 (0.039) | 3.3 |
| S5 | 0.924 (0.031) | 0.926 (0.031) | -0.2 | 0.924 (0.033) | 0.917 (0.035) | 0.8 | 0.918 (0.041) | 0.928 (0.036) | -1.1 |
| S6 | 0.898 (0.052) | 0.919 (0.050) | -2.3 | 0.908 (0.053) | 0.910 (0.062) | -0.2 | 0.916 (0.050) | 0.946 (0.027) | -3.3 |
| S7 | 0.382 (0.096) | 0.340 (0.102) | 11.0 | 0.430 (0.089) | 0.385 (0.098) | 10.5 | 0.556 (0.084) | 0.454 (0.110) | 18.3 |
| S8 | 0.331 (0.095) | 0.268 (0.103) | 19.0 | 0.356 (0.092) | 0.305 (0.102) | 14.3 | 0.498 (0.084) | 0.375 (0.111) | 24.7 |
| S9 | 0.397 (0.108) | 0.252 (0.130) | 36.5 | 0.372 (0.085) | 0.215 (0.087) | 42.2 | 0.531 (0.086) | 0.480 (0.086) | 9.6 |
| S10 | 0.911 (0.032) | 0.910 (0.032) | 0.1 | 0.919 (0.030) | 0.912 (0.028) | 0.8 | 0.956 (0.017) | 0.956 (0.017) | 0.0 |
| S11 | 0.918 (0.038) | 0.918 (0.037) | 0.0 | 0.930 (0.036) | 0.925 (0.030) | 0.5 | 0.963 (0.019) | 0.953 (0.019) | 1.0 |
| S12 | 0.915 (0.053) | 0.916 (0.053) | -0.1 | 0.895 (0.062) | 0.902 (0.051) | -0.8 | 0.958 (0.021) | 0.945 (0.022) | 1.4 |
| S13 | 0.878 (0.033) | 0.879 (0.034) | -0.1 | 0.889 (0.031) | 0.881 (0.035) | 0.9 | 0.920 (0.037) | 0.909 (0.040) | 1.2 |
| S14 | 0.901 (0.030) | 0.909 (0.029) | -0.9 | 0.907 (0.031) | 0.903 (0.032) | 0.4 | 0.933 (0.033) | 0.931 (0.032) | 0.2 |
| S15 | 0.900 (0.036) | 0.892 (0.044) | 0.9 | 0.886 (0.042) | 0.895 (0.042) | -1.0 | 0.937 (0.037) | 0.940 (0.027) | -0.3 |
| S16 | 0.401 (0.094) | 0.346 (0.096) | 13.7 | 0.435 (0.086) | 0.388 (0.089) | 10.8 | 0.579 (0.096) | 0.440 (0.106) | 24.0 |
| S17 | 0.366 (0.095) | 0.309 (0.097) | 15.6 | 0.387 (0.088) | 0.337 (0.093) | 12.9 | 0.538 (0.095) | 0.398 (0.114) | 26.0 |
| S18 | 0.451 (0.122) | 0.330 (0.131) | 26.8 | 0.396 (0.082) | 0.299 (0.071) | 24.5 | 0.597 (0.083) | 0.479 (0.094) | 19.8 |
| **Mean absolute error (MAE)** |  |  |  |  |  |  |  |  |  |
| S1 | 0.115 (0.0197) | 0.114 (0.0180) | 0.9 | 0.102 (0.018) | 0.106 (0.016) | -3.8 | 0.096 (0.016) | 0.103 (0.014) | -7.0 |
| S2 | 0.109 (0.0188) | 0.109 (0.0141) | 0.0 | 0.099 (0.019) | 0.102 (0.015) | -2.7 | 0.097 (0.016) | 0.095 (0.015) | 2.1 |
| S3 | 0.161 (0.0269) | 0.124 (0.0270) | 23.0 | 0.116 (0.029) | 0.118 (0.024) | -1.5 | 0.125 (0.020) | 0.010 (0.020) | 20.1 |
| S4 | 0.119 (0.0115) | 0.113 (0.0157) | 5.0 | 0.1116 (0.012) | 0.1105 (0.015) | 0.5 | 0.086 (0.018) | 0.107 (0.019) | -24.8 |
| S5 | 0.114 (0.0109) | 0.106 (0.0117) | 7.0 | 0.1118 (0.010) | 0.1114 (0.011) | 0.7 | 0.118 (0.010) | 0.104 (0.013) | 12.1 |
| S6 | 0.153 (0.0194) | 0.120 (0.0219) | 21.6 | 0.128 (0.022) | 0.119 (0.025) | 6.7 | 0.139 (0.009) | 0.093 (0.018) | 33.4 |
| S7 | 0.252 (0.0137) | 0.254 (0.0145) | -0.8 | 0.244 (0.013) | 0.246 (0.014) | -0.9 | 0.219 (0.010) | 0.234 (0.013) | -8.3 |
| S8 | 0.259 (0.0197) | 0.265 (0.0223) | -2.3 | 0.255 (0.018) | 0.261 (0.021) | -2.2 | 0.226 (0.010) | 0.247 (0.015) | -9.4 |
| S9 | 0.241 (0.0145) | 0.263 (0.0135) | -9.1 | 0.258 (0.014) | 0.267 (0.022) | -3.4 | 0.220 (0.009) | 0.221 (0.021) | -0.4 |
| S10 | 0.178 (0.0110) | 0.176 (0.0108) | 1.1 | 0.167 (0.011) | 0.165 (0.010) | 1.0 | 0.145 (0.008) | 0.142 (0.008) | 1.8 |
| S11 | 0.146 (0.0128) | 0.143 (0.0123) | 2.1 | 0.136 (0.013) | 0.133 (0.012) | 2.4 | 0.126 (0.010) | 0.127 (0.008) | -0.8 |
| S12 | 0.156 (0.0140) | 0.156 (0.0140) | 0.0 | 0.171 (0.017) | 0.143 (0.016) | 16.1 | 0.125 (0.013) | 0.120 (0.013) | 4.2 |
| S13 | 0.179 (0.0108) | 0.177 (0.0104) | 1.1 | 0.168 (0.010) | 0.172 (0.009) | -2.6 | 0.149 (0.007) | 0.155 (0.007) | -3.5 |
| S14 | 0.151 (0.0112) | 0.145 (0.0110) | 4.0 | 0.146 (0.010) | 0.144 (0.009) | 1.2 | 0.135 (0.006) | 0.133 (0.006) | 1.0 |
| S15 | 0.163 (0.0105) | 0.162 (0.0093) | 0.6 | 0.178 (0.015) | 0.158 (0.014) | 11.1 | 0.141 (0.005) | 0.129 (0.008) | 9.0 |
| S16 | 0.238 (0.0203) | 0.245 (0.0222) | -2.9 | 0.233 (0.018) | 0.238 (0.021) | -2.1 | 0.204 (0.014) | 0.224 (0.018) | -9.9 |
| S17 | 0.249 (0.0200) | 0.253 (0.0240) | -1.6 | 0.246 (0.019) | 0.251 (0.022) | -2.2 | 0.211 (0.014) | 0.231 (0.018) | -9.4 |
| S18 | 0.233 (0.0168) | 0.254 (0.0183) | -9.0 | 0.248 (0.016) | 0.257 (0.021) | -3.6 | 0.203 (0.016) | 0.227 (0.019) | -12.2 |

The three top ranked setups per metric and algorithm choice are highlighted in red (first ranked), orange (second), yellow (third ranked).
The Root Mean Squared Error (RMSE) yielded identical highest rankings, and results highly similar to MAE, data not shown.
For resubstitution tests (*res*), the raw presence data from all $K = 12$ 15°-bands were used to train the SDMs, all else equal to spatial-block CV (*pred*). Both *res* and *pred* are based on 23 analysis scales, with each scale using 1000 sampling folds.

